# Decadal changes in fire frequencies shift tree communities and functional traits

**DOI:** 10.1101/2020.07.22.216226

**Authors:** Adam F. A. Pellegrini, Tyler Refsland, Colin Averill, César Terrer, A. Carla Staver, Dale G. Brockway, Anthony Caprio, Wayne Clatterbuck, Corli Coetsee, James D. Haywood, Sarah E. Hobbie, William A. Hoffmann, John Kush, Tom Lewis, W. Keith Moser, Steven T. Overby, Bill Patterson, Kabir G. Peay, Peter B. Reich, Casey Ryan, Mary Anne S. Sayer, Bryant C. Scharenbroch, Tania Schoennagel, Gabriel R. Smith, Kirsten Stephan, Chris Swanston, Monica G. Turner, J. Morgan Varner, Robert B. Jackson

## Abstract

Global change has resulted in chronic shifts in fire regimes, increasing fire frequency in some regions and decreasing it in others. Predicting the response of ecosystems to changing fire frequencies is challenging because of the multi-decadal timescales over which fire effects emerge and the variability in environmental conditions, fire types, and plant composition across biomes. Here, we address these challenges using surveys of tree communities across 29 sites that experienced multi-decadal alterations in fire frequencies spanning ecosystems and environmental conditions. Relative to unburned plots, more frequently burned plots had lower tree basal area and stem densities that compounded over multiple decades: average fire frequencies reduced basal area by only 4% after 16 years but 57% after 64 years, relative to unburned plots. Fire frequency had the largest effects on basal area in savanna ecosystems and in sites with strong wet seasons. Analyses of tree functional-trait data across North American sites revealed that frequently burned plots had tree communities dominated by species with low biomass nitrogen and phosphorus content and with more efficient nitrogen acquisition through ectomycorrhizal symbioses (rising from 85% to nearly 100%). Our data elucidate the impact of long-term fire regimes on tree community structure and composition, with the magnitude of change depending on climate, vegetation type, and fire history. The effects of widespread changes in fire regimes underway today will manifest in decades to come and have long-term consequences for carbon storage and nutrient cycling.

---

Ecosystem resilience to changing fire regimes^1–3^ will be a key determinant of how terrestrial biomes respond to global change^3–5^. Fire is a pervasive disturbance, burning ~5% of global land area each year and releasing carbon stored in plant biomass equivalent to 20% of anthropogenic fossil fuel emissions^6,7^. Historically, much of this carbon is re-sequestered through time as plants recover and regrow, then lost again in the next fire. However, in many systems, changes in climate and land use have shifted fire frequencies, potentially changing the ability of plants to regrow between fires^1,8–10^.

More frequent burning increases productivity, biodiversity, and plant biomass in some ecosystems, whereas in other ecosystems, little change or even the opposite occurs^11–15^. Our ability to determine why these different responses occur across ecosystems remains limited. At large biogeographic scales, many analyses rely on observational datasets comparing spatial patterns in fire frequency with tree cover and biomass^16,17^; although informative, this approach is limited by the collinearity between variables that both determine fire frequency and tree cover, such as rainfall. Furthermore, the effect of repeated burning on tree cover can take multiple decades to become significant^12,15,18–20^, emphasizing the need to account for the length of time fire frequencies have differed and consider multi-decadal alterations in fire frequencies.

In addition to environmental factors and timescale, plant community composition and species’ functional traits may explain additional variability in responses to long-term changes in fire frequency^21^. For example, traits related to physiological protection from heating during fire and the capacity to colonize and regrow rapidly could help predict losses of trees due to frequent burning^15,22–24^. Additionally, nutrient acquisition and use traits can influence the future productivity of plants and their ability to regrow after fire^25^ and also have long-term implications for carbon and nutrient cycling in soils^26^. For example, plants that form symbioses with ectomycorrhizal fungi, arbuscular mycorrhizal fungi, or nitrogen-fixing bacteria may be better equipped to access limiting nutrients under frequent burning. The distinction between strategies is important, however, because ectomycorrhizal plants tend to slow nutrient cycling and productivity, while arbuscular and nitrogen-fixing species can accelerate cycling and increase productivity^25–28^.

The existence of experimental manipulations of fire frequencies across sites that span large environmental and compositional gradients offers an opportunity to test how ecosystems respond to altered fire frequencies^29–32^. However, the few studies comparing multi-decadal fire manipulations have typically been constrained to small groups of sites within single ecosystems^31,33^. Here, we quantified the effects of fire frequency on tree cover across broad biogeographic and climatic scales, incorporating additional factors that may explain variability in fire effects on tree communities. We analyzed data on tree populations from 29 sites and 374 plots; at 27 of the sites (324 plots), surface fire frequency was experimentally manipulated for 16-64 years (mean of 30 years), and at two sites (50 plots), natural variation in crown fire frequency presented a natural experiment. The sites cover North and South America, Africa, and Australia across major biomes that experience frequent burning (Figure S1, Table S1, *Supplemental Information, SI*). Each surface fire site contains replicate plots including an unburned treatment and different prescribed burning frequencies (Figure S2), where fire frequencies ranged from approximately one fire every decade to one fire every year (Table S1). To evaluate the effects of fire alone and in combination with environmental covariates while accounting for the high variability in overall tree basal area and stem density across sites, we used mixed-effects models with site as a random intercept as the main test in our analysis (*SI*). We focused on tree responses because trees are critical for long-term carbon storage and productivity^34^, define the ecosystem (e.g., whether a landscape is a forest or savanna)^35^, and influence several biogeochemical processes^36^.

There was a clear overall effect of fire treatment on tree populations. Tree density (stems per-hectare) tended to be lower in frequently burned plots relative to infrequently or unburned plots (Figure 1a). A comparison between the most extreme fire frequency treatments using response ratios illustrated that densities were 44±25% lower in the most frequently burned plots compared with unburned plots, and that the differences between fire treatments were lower when the differences in fire frequencies were lower (Figure 1b, Table S2, error defining 95% confidence intervals). When fire frequency and duration of study were analyzed as continuous variables across all plots and sites using mixed-effects models, sites with longer durations of altered fire frequencies had larger differences between fire treatments (F_1,282_=47, p<0.001), with the slope between duration and stem density being more negative the more frequently plots were burned (F_1,280_=8.4, p=0.004, Figure 1c). For example, relative to unburned plots, stem density in plots with a three-year fire-return interval was 26% lower after 30 years and 48% lower after 50 years (Figure 1c, Table S3). Fifty years of annual burning resulted in burned plots having 63% lower stem density relative to unburned plots (Figure 1c, Table S3, see Figure S3 for non-transformed results).

**Figure 1:**
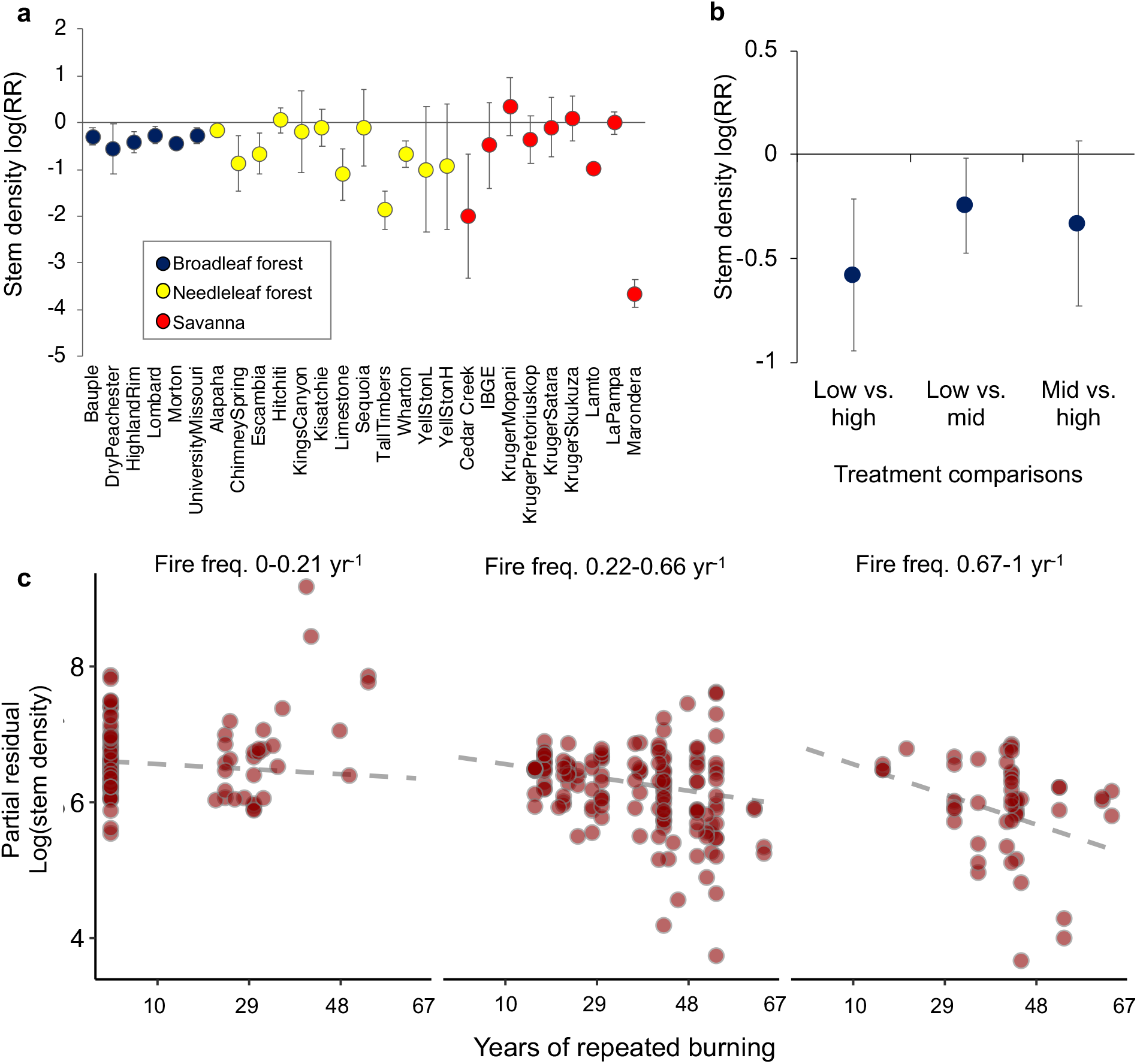
Fire effects on stem density increase with degree of frequency contrast and length of study duration. **a-b**) log response ratios of stem densities and the surrounding 95% confidence intervals. **a**) comparisons within each individual site colored by broad biome categorization in most extreme fire frequency treatments loge(burned/unburned) (Table S1). **b)** comparisons among the different levels of fire frequency in studies with ≥3 levels; less frequent treatment always in denominator, Table S2). **c**) partial residuals plot from a mixed effects model including fire frequency, the number of years of repeated burning, and their interaction for loge stem density (Table S3); site was used as a random intercept. Panels are centered on cross-section values of one fire every 10 years, 1 every 3 years, and 1 every year but encompass a range of fire frequencies within each panel.

Fire type was also important, with frequent crown fires affecting tree populations to a greater degree than frequent surface fires. Comparison of 50 plots in needleleaf forests that experienced natural variability in the frequency of stand-replacing crown fires (i.e., wildfires) illustrated that stands with shorter fire-return intervals had significantly lower tree densities, especially when plots with the shortest return intervals were considered (F_1,26.5_=5.2, p=0.03 and F_1,21_=10.3, p=0.004, Figure S4). Experimental manipulation of surface fire frequency (i.e., prescribed fires) in needleleaf forests in the USA showed that stem densities were lower in more frequently burned plots, but less so than differences caused by frequent crown fires (F_1,47.1_=17.2, p=0.001, Figure S4). The large effect of short-interval crown fires on tree communities, supported by studies from other regions^37,38^, highlights the importance of higher fire intensities having more severe effects.

Fire had similar effects on tree basal area, which we analyze in detail, because basal area correlates with tree biomass, canopy cover, and tree carbon storage. Basal area was on average 54±25% lower in the most frequently burned plots relative to the unburned plots (Figure 2a,b). When frequency and duration were considered in parallel across all sites, the lower basal area in frequently burned plots became more apparent with increasing experimental duration and frequency of burns (frequency-duration interaction, F_1,289_=23.3, p<0.001; Figures 2c, Table S3, mixed-effects models). For example, plots with 30 years of triennial burning had 27% less basal area relative to unburned plots, while those with 50 years had 53% less basal area. Divergence between fire treatments was even greater after 50 years of the most extreme frequencies of annual burning, where burned plots had 72% less basal area than unburned plots. Consequently, changing fire frequency and duration of exposure shifted tree basal area and stem abundance across sites. The manifestation of fire effects began to lessen as experiments had increasingly long durations, suggesting that effects will saturate as tree cover approaches a new equilibrium.

**Figure 2:**
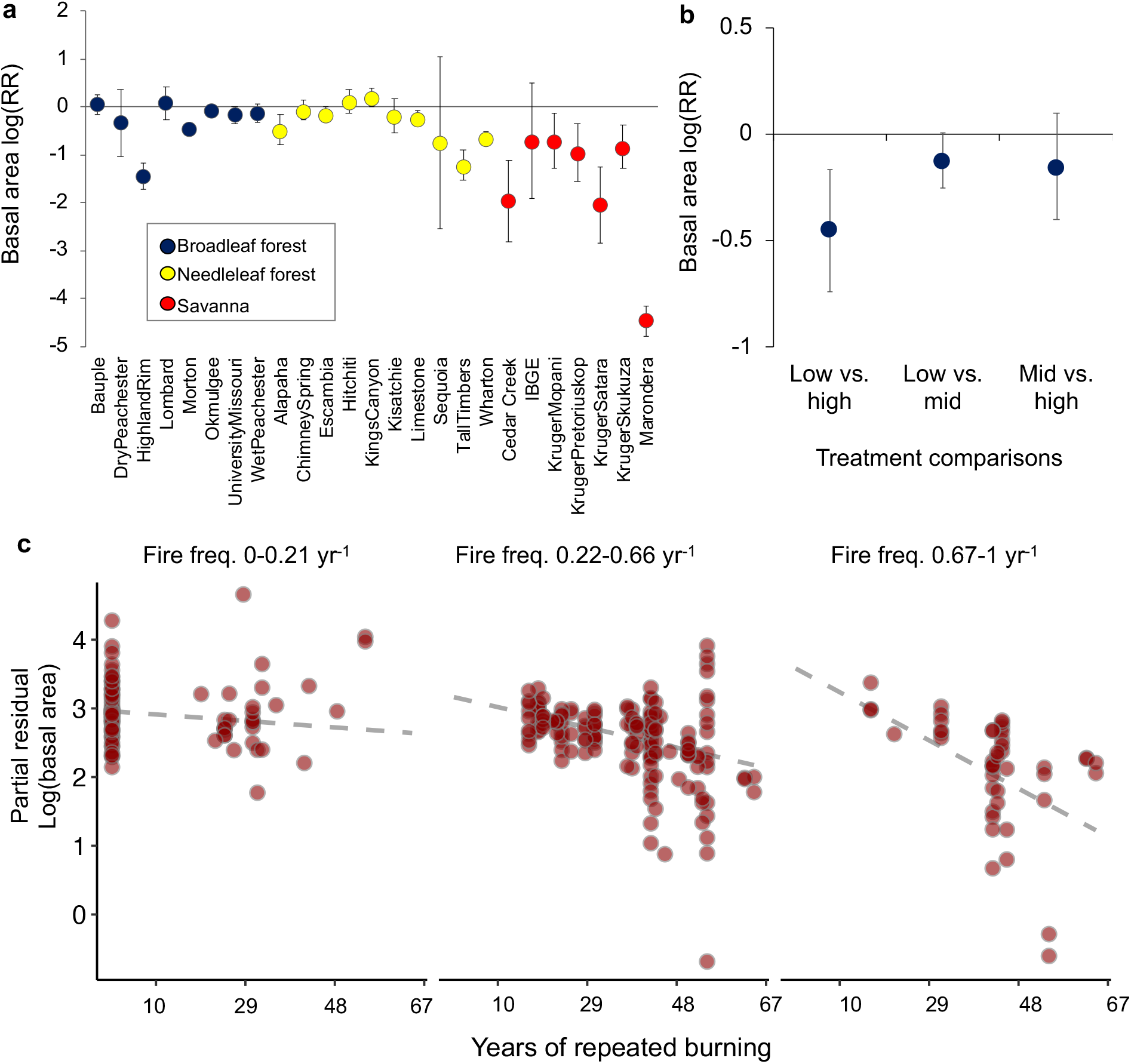
Frequent burning decreases tree basal area and compounds with time of exposure to different fire frequencies. **a-b**) log response ratios of basal area and the surrounding 95% confidence intervals. **a**) comparisons within each individual site colored by broad biome categorization in most extreme fire frequency treatments loge(burned/unburned) (Table S1). **b)** comparisons among the different levels of fire frequency in studies with ≥3 levels; less frequent treatment always in denominator, Table S2). **c**) partial residuals plot from a mixed effects model including fire frequency, the number of years of repeated burning, and their interaction for loge basal area (Table S3); site was used as a random intercept. Panels are centered on cross-section values of one fire every 10 years, 1 every 3 years, and 1 every year but encompass a range of fire frequencies within each panel.

The effects of changing fire frequencies also depended on the fire history of the site prior to the establishment of the experiment. In forest sites that burned regularly in the decades prior to the onset of the experiment, fire exclusion resulted in basal area being 50% (±17%) higher than treatments that maintained historical burning frequencies (p=0.002, Figure S5, Table S1 for site fire histories). In contrast, the reintroduction of fire into forests that had not burned for several decades prior to the onset of the experiment had relatively minimal effects (p=0.13, Figure S5). These results differ from studies on wildfires which are known to have larger effects in forests that have a history of fire exclusion due to high fuel accumulation^39,40^, which is somewhat expected given the lower severity of prescribed surface fires. In savannas, where the fire experiments were all initiated in landscapes that burned regularly in the decades preceding the experiment, fire exclusion resulted in basal area increasing by 41% (±20%), but increasing fire frequency resulted in basal area declining by 48% (±16%), relative to an intermediate interval that maintained the pre-experiment frequency (statistics from log response ratios ±95% confidence intervals, p<0.001 for both, Figure S5, *SI*). Taken together, the largest effects of altered fire frequencies were due to fire exclusion in landscapes that had burned regularly for at least the past few decades.

Climate played an important role in modifying the effect of fire frequency on trees. Fire effects were largest in areas that received more rainfall in the wet season, less rainfall in the dry season, and had lower mean annual temperatures (F_1,292.2_=55.2, p<0.001, F_1,284.7_=9.8, p=0.002, and F_1,283.2_=18.1, p<0.001, respectively) (Figure 3a-c, Table S4, see Table S5 for stem density). Sites with higher precipitation in the wet season experienced larger effects of burning. For example, plots that experienced more frequent burning (2 fires every 3 years, one standard deviation above mean frequency) had 67% lower tree basal area than unburned plots in sites with high wet season precipitation (Figure 3a, S6, Table S4, see *SI* for details on calculations). The difference between treatments was only 22% in sites with average wet season precipitation.

**Figure 3:**
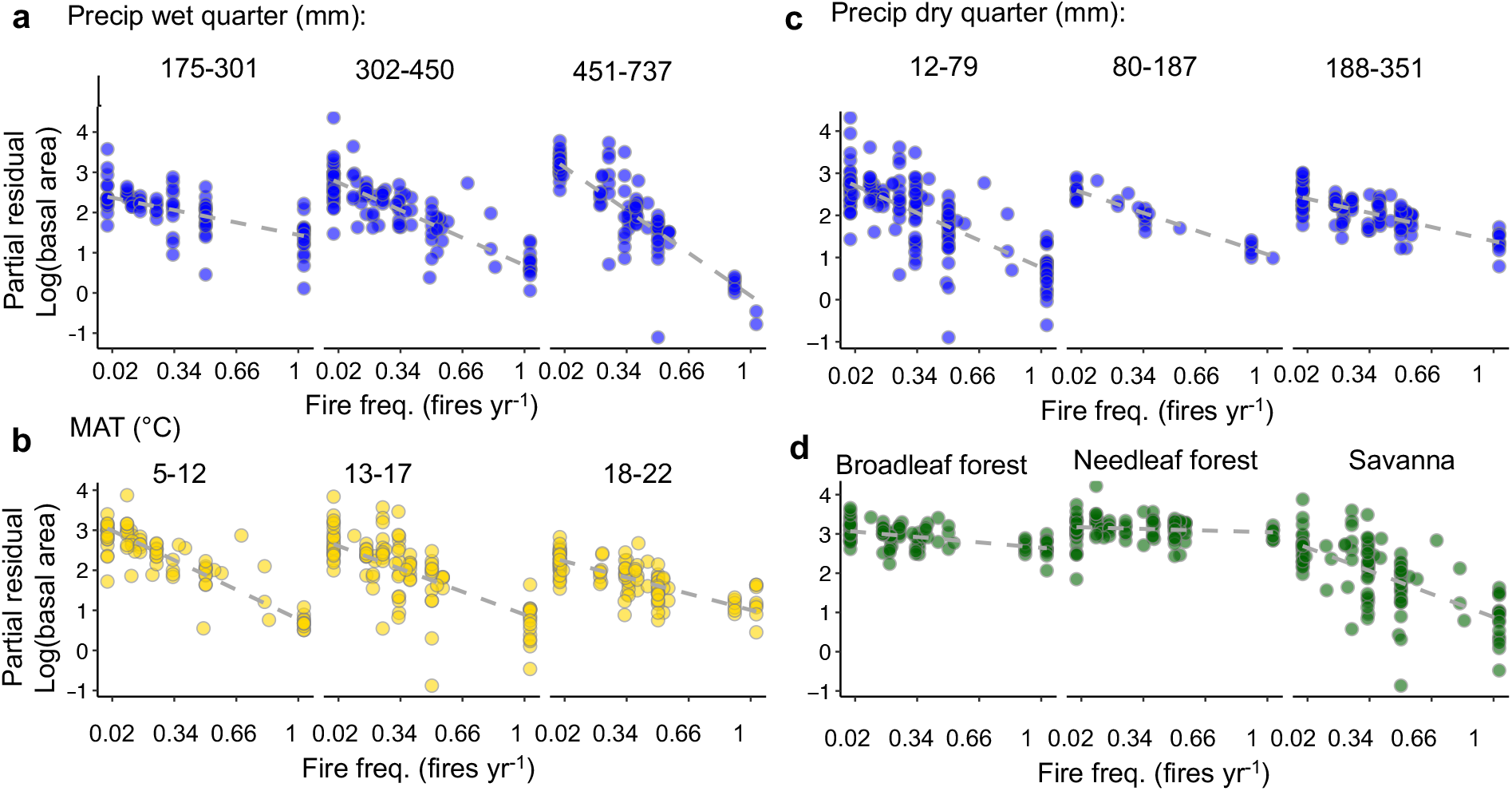
Climate, ecosystem type, and plant traits modify effects of fire frequencies on tree basal area. Partial residual plots of the mixed-effects model illustrating the interactive effects between covariates (site as a random intercept). Panels centered on cross-sectional values from one standard deviations around the median (−1, 0, 1). MAT: mean annual temperature. Comparisons of rainfall scenarios relative to the mean in the text used wet-season precipitation of +1 standard deviation above the mean (525 vs. 375 mm yr^-1^) and dry season precipitation of −1 standard deviation below the mean (25 vs. 133 mm yr^-1^). The duration of experiment held at its mean of 28 years. All model fits are p<0.05; statistics are in Table S4.

Precipitation in the dry season had opposite effects. Sites with lower precipitation in the dry season experienced twice as large an effect of fire on basal area (46% vs. 22% lower tree basal area in sites with low vs. average dry season precipitation Figure 3c, Table S4). The contrasting response to precipitation in the wet vs. dry season is consistent with our understanding that fires are most intense in areas with stronger wet seasons (leading to more fuel) and more severe dry seasons (lower fuel moisture), thus contributing to potential losses with more frequent burning^41,42^. Rainfall in the dry season likely also influences fire effects by determining the water available for tree growth when fire is excluded. Soil characteristics did not explain sensitivity to changing fire frequencies across sites; across all sites, neither texture-based classification of soils nor soil carbon content interacted with fire frequency (Table S4).

The effect of fire on tree basal area also differed across ecosystems (F_2,279_=14.5, p<0.001), with frequent burning having a larger effect on tree basal area in savannas relative to broadleaf and needleleaf forests (accounting for climate effects and differences among continents, Figure 3d, Table S4). Relative to the unburned plots, basal area in frequently burned plots was 6% lower in needleleaf forests and 22% lower in broadleaf forests (Figure 3d, burn frequency of two fires every three years, *SI*). In savannas, frequently burned plots had 70% lower basal area relative to the unburned plots (Figure 3d, Table S4). Interestingly, stem density responses to fire frequency were qualitatively different between savannas and forests (Table S5). Stem densities increased with more frequent burning in forests while basal area decreased, potentially due to higher light availability and recruitment of trees in the forests. We tested the sensitivity of our findings that savannas were more sensitive to increased burning frequency via a subdivided classification of ecosystems by partitioning broadleaf forests into oak and eucalypt types and needleleaf forests into those that transitioned between oak and pine dominated (Table S1). When included in the final model, the subdivided vegetation classification still had a significant main effect (F_4,19.4_=12.4, p<0.0001), and a significant interaction with fire frequency (F_4,276.8_=7.8, p<0.001, Figure S8), with basal area in savannas responding the most to changes in fire frequency (Figure S8).

We next tested the extent to which plant traits influence tree responses to fire across ecosystems^23^. We analyzed only the experiments from North America (77 tree species, 16 sites, 181 plots) because trait data were available there to (*i*) categorize species by nutrient-acquisition strategies, and (*ii*) assign wood, leaf, and root traits related to growth, survival, and nutrient-use strategies. Plots with tree species having thinner bark and denser wood changed relatively more with frequent burning (bark: F_1,154.3_=5.7, p=0.018; wood density: F_1,154.1_=12.9, p<0.001, Table S6, Figure 4a,b). Within a site, mean wood density of the tree community tended to be lower in frequently burned landscapes, potentially because of increasing dominance of gymnosperms, which tend to have lower wood density. In contrast, we did not observe any effect of fire on the mean bark investment of the tree community (Table S6), demonstrating that bark investment at the community scale does not appear to change in response to fire; nevertheless, bark investment may influence basal area loss patterns across broad biogeographic scales.

**Figure 4:**
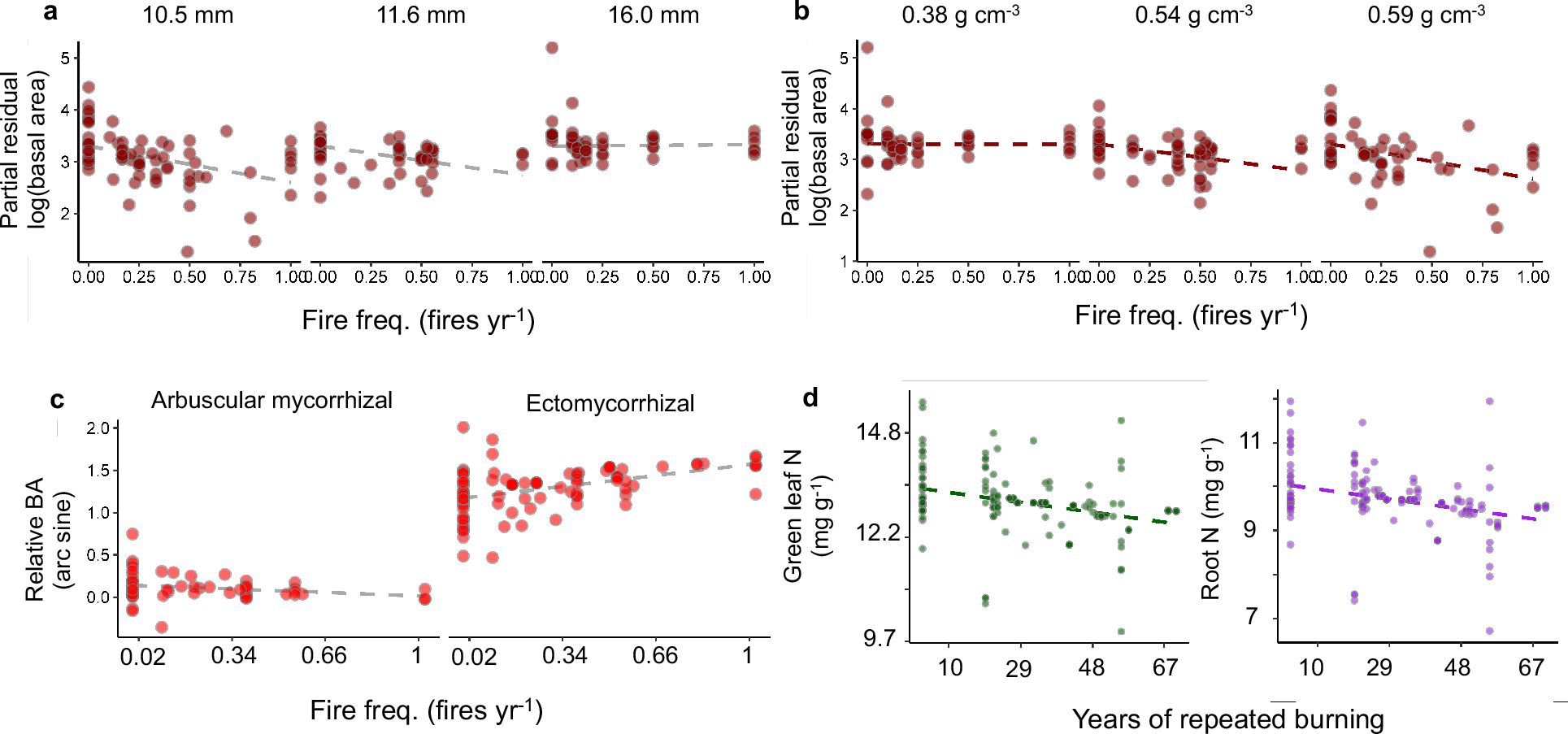
Frequent burning filters for conservative nutrient-use and acquisition strategies across North America. Partial regression plots from mixed-effects models with community weighted means of (**a**) bark investment scaled to a 10 cm stem size and (**b**) wood density (WD) as modifying variables. Bark and wood density were included in the same model and were negatively correlated σ = −0.86. Statistics are in Table S6. Basal area is loge transformed. **c-d**) community weighted means of nutrient acquisition (c) and nutrient use (**d**) strategies. c) relative basal area (arc sine transformed) of arbuscular vs. ectomycorrhizal trees as a function of fire frequency. **d**) green leaf and live root N as a function of study duration. Statistics are in Tables S7–S9. Data on litter N and resorption are in the Tables and phosphorus content data are in Figure S8.

Frequent burning also shifted the nutrient use and acquisition strategies of tree communities, as expected given the nitrogen (N) losses resulting from frequent burning^43^. Plots burned frequently for longer periods of time were dominated by tree species with low N concentrations in green and senesced leaves and roots, and resorbed a greater proportion of N before leaf senescence (p<0.001 for all variables, Figure 4d, Table S7). Tissue phosphorus (P) concentrations also declined with frequent burning in leaves and litter but not in roots (Figure S10, Table S7).

Fire also affected the relative abundance of nutrient-acquisition strategies. We evaluated changes in acquisition strategies using categories of trees’ abilities to form symbioses, which correlate with several other traits involved in acquisition^44^. Trees that formed symbioses with ectomycorrhizal (ECM) and arbuscular mycorrhizal (AM) fungi were the most abundant nutrient-acquisition strategies across our plots; ericoid and nitrogen-fixing trees were absent from most sites (Figure 4c). ECM trees, which contain fungal symbionts capable of acquiring N from organic matter^45^, tended to be more successful in frequently burned plots. The relative abundance of ECM trees increased from 85% in unburned plots to nearly 100% in annually burned plots (Figures 4c, Table S8). ECM trees were also more common in warmer climates and on soils with low organic matter (Figure S11, Table S8). ECM trees typically have lower concentrations of N and P in leaves, litter, and roots than AM trees^46^ (Figure S12, Table S9), suggesting the turnover in symbiont composition may be driving the shift in stoichiometry of the tree community. The tendency for frequently burned plots to have tree communities dominated by ECM trees with low N and P content in leaves, roots, and litter indicates that frequent burning favors tree species with a suite of traits consistent with a conservative strategy of nutrient use and acquisition.

Although our analysis is to our knowledge the largest compilation of results from fire manipulation plots to date, it identifies several factors that highlight the need for even larger-scale analyses. For one, an improved representation of fire experiments in different ecosystem types across continents (e.g., tropical forests in Africa and savannas in Australia) will help further unpack the variability across ecosystems. Past research has demonstrated that the turnover in tree species composition can be important for explaining changes in total tree cover within experiments^47^. Thus, a better understanding of how fire kills trees and how these processes differ across ecosystems may give a more complete picture of why fire effects vary. Our analyses also demonstrated that land use history and the fire regime prior to experiment establishment are critical to interpreting the magnitude of fire effects, consistent with previous studies^48^. Consequently, global-scale estimates of how current shifts in fire frequency alter ecosystem carbon should carefully consider how uncertainties in fire history preceding the satellite era may influence their estimates.

Our findings that fire effects emerge over multiple decades but then approach a new (non-zero) equilibrium are in agreement with studies that have performed repeated measurements of tree populations within the same experiment. Generally the treatments can diverge over the first few decades^15^, but the rate of divergence declines through time^30^. The timescale of change is similar to the multi-decadal shifts in soil carbon and nitrogen, which likely reflect a link between tree biomass inputs into soil carbon pools and potentially the turnover of plant traits interacting with changes in soil nitrogen pools^25,49,50^.

In conclusion, widespread changes in fire regimes are likely to have structural, compositional, and functional effects on tree communities that manifest over decades. Importantly, fire is an integral part of many ecosystems and can promote biodiversity, reduce wildfire risk, and stimulate nutrient turnover; consequently, lower tree basal area and density in more frequently burned plots is not necessarily a negative result depending on the management goals for the ecosystem. Nevertheless, persistent changes in fire frequency will have, and already are having, profound effects on ecosystems and need to be considered in projections of communities and ecosystems in the future.

## Supplemental Information

### Experimental design and site descriptions

The majority of sites sampled comprise ecosystems that experience surface fires (from fire manipulation experiments, n=27). Our main analyses are based on the surface fire experiments, but we compare these data with a network of plots across n=2 sites with natural variability in the frequency of stand-replacing crown fires to evaluate the effect of fire regime. We describe the sites briefly in Table S1, and present detailed descriptions of site history in Dataset 1.

The surface fire experiments mostly are experimental prescribed burn plots. The managers generally try to burn in a broad seasonal window (e.g., a spring fire in North America may occur anytime from March-May) to optimize burn timing for the local fire conditions most suitable to their planned fire intensities. The sites contained different land use histories before the establishment of the experiment, which was not always documented in detail, but we describe key factors in Tables S1 and Dataset 1. We describe how we evaluated the potential role of land use history in *Testing role of fire and land use history* below.

Because these experimental sites utilized different survey methods, the classification of a plant as a tree differed. In some cases, such as savannas with relatively small woody plants, all woody plants over a basal diameter of 5 cm were measured (which includes shrubs). In other cases, stems only above 10 cm diameter at breast height were measured. Consequently, the definition of ‘tree’ is based on the local knowledge of what is the relevant size threshold for a particular ecosystem, and in some cases it includes all woody plants.

The stand-replacing crown fires are from one extensive ecosystem type that accounts for a large amount of forest fire area in North American temperate regions. Specifically, we used data from 50 plots in lodgepole pine forest in the Western United States^23^ (n=50) spanning different elevations and plots along a continuum of fire return intervals^23^. Because this ecosystem experiences stand-replacing fires, time-since-fire is critical for determining tree abundance because it determines the stage of regrowth. We dealt with this by sampling plots that differed in their fire return interval over the past several hundred years but shared the same time since last fire because of a large fire that burned forests with different times since the previous fire. Given the previous study found elevation to be important, we included elevation categories in the model (<2400 m and >2400 m, respectively).

### Choice of plots within 27 sites with surface fires

Within each site, we only used one sampling time period for our analysis. Eight sites contained time series data: Cedar Creek, Lombard, Sequoia, Kings Canyon, and the four Kruger sites. For Lombard, we used the surveys from 2002, for Cedar Creek we used 2010, for Kruger we used 1996-98 and for Sequoia and Kings Canyon it varied according to the replicate plots because their most recent surveys occurred in different years. For Cedar Creek, more recent surveys exist, but the outbreak of oak wilt has resulted in large amounts of tree mortality not due to fire (Reich *personal communication*). For Highland Rim, we used two different sets of data: the first dataset contains plot-level data (thereby allowing us to determine a variance around the mean) but no tree species identities; the second dataset contains no plot-level data but has treatment-level averages within each tree species, which allowed us to analyze composition changes. We utilized the plot-level data for analyses of basal area and stem density. For Morton and Okmulgee, there are not always true replicates in each fire treatment. Morton contains two true replicates for the unburned, but no true replicates for the burned plots. Okmulgee contains no true replicates.

In sites where fires were prescribed in different seasons, we used a single burn season in the analysis in an attempt to match seasons of burns within a particular ecosystem type within a particular region. In North America, we standardized fire season to burning conducted in the winter to early spring because not all sites contained fire treatments with summer burns. For Hitchiti, we used the December-March burns, dropping the June burns. For Kisatchie we used March, dropping July and May. For Kruger, we used August, dropping all the other seasons. For Lombard, we used March-May, dropping the June-Aug.

Lamto and La Pampa only contained data on the number of tree stems. Consequently, these were incorporated into the stem abundance analysis only.

### Soil chemistry data

We collected and analyzed soil data using several methods. First, we determined the dominant soil type using either author descriptions or reported soil texture analysis. Second, we used the highest resolution soil data as possible (e.g., soil samples from each replicate plot within a fire treatment), but some sites only contained site-level soil properties. Consequently, we analyze overall effects of fire on all sites without any covariates, followed by a model that uses model selection to account for collinearities among variables when testing for factors that modify fire effects. To extend data on soils across plots, we sampled soils (top 0-5 cm of the mineral horizon) in 24 plots across four sites: Kings Canyon, Sequoia, Limestone Flats, and Chimney Springs. Each site contained three replicate plots of an unburned treatment and a high fire frequency treatment. We collected n=5 pseudo-replicates within the true replicate plot, analyzed the soils for carbon, nitrogen, and texture, and averaged within each plot.

### Climate data

To obtain long-term climate averages at each site, we used WorldClim^51^. Managers timed burning to coincide with consistent weather conditions over the course of the experiment, therefore we did not obtain high resolution inter-annual variability in climate. We focused on several climate variables based on ecologically relevant *a priori* hypotheses: (i) precipitation partitioned into the driest and wettest quarters of the year because precipitation influences fuel accumulation (primarilly in the wettest quarter) and fire conditions (primarily in the driest quarter) and (ii) mean annual temperature because of its large effect on a variety of biogeochemical processes. Precipitation in wet and dry quarters are not as correlated with one another but are highly correlated with mean annual precipitation and temperature (Table S10).

### Calculation of fire effects in different environmental conditions

Several methods exist to calculate variable importance, with no clear optimal method ^52^. We chose to use the regression coefficients in the model to understand the sensitivity of basal area and stem density to changes in relative values of each variable. Importantly, the models were fit to re-scaled data by subtracting each value by the mean and dividing by the standard deviation of the variable. Consequently, the product between the mean value of a variable and its coefficient is always zero. Thus, we can compare the relative impact of variables by comparing the magnitude of the fitted coefficients because they reflect the potential change in basal area for a one standard deviation change in a variable value.

To perform meaningful comparisons, we use the standard deviations of variables to illustrate the sensitivity of basal area to a change in the value. For example, using the model to estimate the effect of increasing fire by 1 standard deviation from the mean (mean = 0.34, mean + 1σ = 0.67) tells us the sensitivity of basal area to fire, with all other variables held at their means. Interactions can be tested by moving two variables away from their means: for example, changing the fire value in conjunction with precipitation in the wet quarter. Because the model is fit to re-scaled data, the intercept of the model is not representative of the unburned fire treatment, which is calculated by re-scaling the fire frequency data (0-μ)/σ, which gives a value of −1.081, making the unburned calculation of 25.6 m^2^ ha^-1^ when all other variables are held at their means.

Here are the different levels of comparisons we used in the results and the corresponding figures.

*Wet season precipitation* (Figure 3a): wet season precipitation varied one standard deviation above the mean vs. at the mean (525 vs. 375 mm yr^-1^, respectively). Fire frequency varied from unburned to one standard deviation above the mean (2 fires every 3 years).

*Dry season precipitation* (Figure 3c): dry-season precipitation was one standard deviation below the mean vs. at the mean (25 vs. 133 mm yr^-1^, respectively). Fire frequency varied from unburned to one standard deviation above the mean (2 fires every 3 years).

*Vegetation type* (Figure 3d): fire frequency effects were made using two levels of comparisons. Unburned plots vs. burning at the mean frequency (1 fire every 3 years) and unburned plots vs. burning at one standard deviation above the mean frequency (2 fires every 3 years).

### Testing overall fire effects

We first tested the overall effects of the fire treatments across sites with log response ratios using techniques employed meta-analyses^53,54^. First, we calculated the log response ratio between the different fire frequency categories (low, medium, and high) for basal area and stem density averaged within each category, with the lowest fire frequency in the comparison always in the denominator. Next, we determined the variance based on the number of true replicates within each treatment in a site and the standard deviations within the fire frequency category. These values across sites were then used to determine the effects of fire treatments on tree basal area and stem density.

We first evaluated the overall effect of fire frequency and length of time frequency was altered on tree basal area and stem density without considering any potential modifying role of covariates to test the general effect of fire across all sites. To accomplish this we analyzed (i) a mixed-effects model containing fire frequency, fire period, and their interaction, and (ii) log response ratios of stem density and basal area relativized within each site. We excluded the 50 crown fire plots for this initial analysis. We fit the mixed-effects models with site as a random intercept. The statistical design is nested because each site has several replicate plots receiving different fire treatments. As a result of this design, the responses to fire at the plot level are likely more related within sites than between sites, necessitating a random intercept. Although our design is not balanced (sites differ in their number of replicate plots), models are generally robust to unbalanced designs unless sample sizes are low and/or a random slope is being estimated ^52^, neither of which are applicable here. Models were constructed based on our *a priori* hypotheses of how fire would influence tree population sizes and the potential to interact with covariates. In all cases of mixed-effects models, we tested for model significance using Satterwaith’s approximation for degrees of freedom and a Type III ANOVA ^55^. In the event of an insignificant main effect but significant interaction, we tested whether the main effect could be dropped from the model using a change in Aikake Information Criterion (AIC) with a threshold of two.

### Comparison between surface vs. crown fire regimes

To analyze the effect of crown vs. surface fire types, we analyzed stem density data from 50 plots (paired within 25 locations) in the Western USA in a separate model. All plots had the same time since fire of 12 years. For this analysis, we used a mixed-effects model to test the relationship between fire return interval and stem density for all locations across the entire return interval span with location as a random intercept. As a further test of fire return interval effects, we selected the short fire return interval (<100 years) in each paired plot and analyzed the relationship with a linear model.

### Testing the role of fire and land use history

We partitioned studies into three categories based on their disturbance history. Using knowledge of fire history for several decades prior to the fire experiments, we determined if the fire treatments within a site reflected (i) an increase in fire frequency above a historical mean, (ii) fire exclusion after decades of repeated burning prior to the experiment, and (iii) reintroduced fire after decades of pre-experiment fire exclusion (Table S1); the historical mean was defined based on fire activity data for several decades prior to initialization of the experiment (Dataset 1).

The fire experiments in the savannas were all initiated in sites that had regularly burned for several decades before the establishment of the experiment. The intermediate fire frequency treatments were reflective of the historical mean, but the most frequently burned plots in those sites were burned at a frequency higher than the historical mean. Consequently, we could use the intermediate frequency plots to evaluate the relative changes due to fire exclusion (unburned vs. intermediate) or tree cover declines because of more frequent burning (frequent vs. intermediate). In one savanna site, Marondera, all trees were removed before the onset of the experiment, and consequently we are not able to assume that the difference between the intermediate and high frequency treatment is due to declines in trees since the onset of the experiment, rather, it is likely due to a restriction on recovery. Consequently, we omit Marondera from these calculations.

The fire experiments in the forests varied in their historical fire frequency and the occurrence of other disturbances. Several sites were in some stage of recovery from previous land use (e.g., selective logging, agriculture, etc.), but we focused on the variability in fire history to categorize the sites into fire response categories. We partitioned forests into those that had remained unburned for several decades before the onset of the fire treatments (i.e., reintroduction burns) vs. sites that burned regularly before the experiment. We assume that in the case of the reintroduction burns, changes in tree cover arises from losses due to more frequent burning.

In the sites that burned regularly prior to the establishment of the experiment, the differences between the unburned plots and those burned at the historical mean was assumed to arise from gains under fire exclusion, and not necessarily increased losses due to frequent burning (although that can clearly occur).

We analyze the effect sizes of fire in the comparisons of the unburned vs. intermediate vs. frequent treatments using the same meta-analysis method described above.

### Model selection to determine parsimonious variable combinations

For the plots with surface fires, we performed model selection by incorporating covariates of climate, soil, and plant composition into mixed-effects models to test for pairwise interactions and possible collinearities (see discussion below of collinearities). Finally, we constructed a full model containing fire, climate, soil, and composition variables based on our hypotheses that these factors will interact with fire frequency as well as information gained from the pairwise tests. There were several insignificant effects in the final model, which we tested for removal using model selection with a threshold AIC of two. All variables were re-scaled by subtracting the mean and dividing by their standard deviation.

Our selection process in the tables illustrates the sensitivity of the final model to the inclusion of additional interactive effects that are not in the final model as well as main effects of the climate, geography, and soil variables. We do not present the exhaustive comparisons because they are not guided by our *a priori* hypotheses of factors modifying fire effects. Soil type was not reported for one location with stem density measurements in South America, so we just use soil carbon content in the model selection analysis.

### Evaluating assumptions of aggregating ecosystem types

The vegetation composition at each site differs substantially, ranging from diverse tropical savannas with dozens of tree species (e.g., Kruger sites) to monodominant coniferous forests (e.g., Limestone Flats and Chimney Springs). Classifying the sites into broad categories was done methodologically, by balancing the need to maintain parsimony (and thus statistical power) with accurately capturing how plant composition may modify fire effects. Consequently, we performed two levels of classification: (i) a coarse categorization based on biomes, as savannas vs. forests, and within forests treating broadleaf and needleleaf forests separately, which we refer to as a vegetation type; and (ii) accounting for variability within forest types by partitioning broadleaf forests into Myrtaceae (eucalypt) vs. Fagaceae (oak) dominated, and needleleaf forests into forests that are near completely dominated by needleleaf trees vs. a mixed forest containing both needleleaf and broadleaf trees, which we refer to as a sub-vegetation type.

### Collinearity among climate variables

Climate variables can be highly collinear, which can inflate the risk of error in statistical inference. To evaluate collinearity, we first determined the Pearson correlation coefficients between the main climate variables. We excluded variable combinations with a correlation >0.70. Most climate variables relating to water availability were not correlated with mean annual temperature. For water availability, we used precipitation in the driest quarter and the wettest quarter because their correlation coefficient was relatively low and they are ecologically more relevant than annual means because they determine the potential productivity in the wet season when most growth occurs but also potential water stress and fire conditions in the dry season (Table S10). In contrast, mean annual precipitation and aridity were tightly correlated with one another, as well as with the precipitation values in the separate quarters.

### Species classifications and functional traits

Bark thickness data were collected from a dataset in the Fire and Fuels Extension of the Forest Vegetation Simulator. https://www.fs.fed.us/fmsc/ftp/fvs/docs/gtr/FFEaddendum.pdf. Although broad syntheses of bark investment exist for many tree species in North America, not all species contained data from empirical measurements, and thus we used the data from the Fire and Fuels Extension. Bark thickness was assumed to scale linearly with stem diameter, which is generally valid for smaller stems, but it is known bark saturates with increasing stem diameter ^56^. The ability of bark investment to predict fire effects will likely improve with better consideration of the non-linear relationship between bark and stem diameter. We evaluate the relative bark investment, and not absolute bark thickness, which is based on bark investment as well as stem size.

Wood density was compiled from the literature using a global wood density database ^57^, supplemented with additional data ^58,59^. We assigned a genus-level average for 19 species lacking data.

Plant tissue stoichiometry and mycorrhizal type were determined using both trait data as well as phylogenetic trait estimates calibrated to trait data used in a previous global analysis of plant mycorrhizal traits ^46^. Full data selection criteria are presented in ^46^, but we describe them briefly below.

The plant phylogeny contained >49,000 plant species ^60^. Plant species were added to this phylogeny as needed using the *congeneric.merge* method ^61^. This method uses congeners to add species missing genetic data to the phylogeny, conservatively replacing genera with polytomies where more than one member of the genus is present in the analysis.

We next generated a species-level phylogenetically estimated trait value for each species and trait by fitting models to all data for a particular trait as a function of phylogenetic distance, leaving out each species one at a time using the phyEstimate function within the picante package for R statistical software^62^. This way, each species trait estimate is based on its own phylogenetic position and a phylogenetic model of evolution (Brownian motion) parameterized without that specific species trait observation. For species without trait data, we estimated trait values based on a model fit to all available trait data.

### Testing the interactions between species composition and fire

To test for fire effects on the relative abundance of symbiotic strategies, we calculated the relative basal area of the different strategies (ectomycorrhizal, arbuscular mycorrhizal and the less abundant ericoid mycorrhizal, non-mycorrhizal, and nitrogen-fixing tree species). Given the low occurrences of ericoid, non-mycorrhizal, and nitrogen-fixing species, we analyzed the relative abundance of arbuscular mycorrhizal and ectomycorrhizal species only. We then fit mixed-effects models with relative basal area as the dependent variable and fire, climate, broad vegetation type (broadleaf, needleleaf, savanna), and soil conditions as the independent variables, each modified by a symbiont term. Relative basal area was arcsine transformed. This analysis was conducted in the North American plots.

To test how functional traits correlated with the effects of fire frequency and duration of experiment, we calculated community trait means in plot *j* by averaging the traits of each species *i* by their relative basal area (BA) in a plot. Bark thickness (Bark) for example:

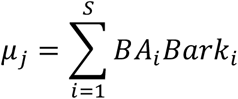

We calculated community weighted means (CWM) for wood density, bark thickness, live and senesced leaf nitrogen (N) and phosphorus (P) and live root N and P. We also calculated retranslocation of N and P from a live leaf before senescence using the data from live and senesced leaf N and P (i.e., not directly measured). Calculations using N as an example:

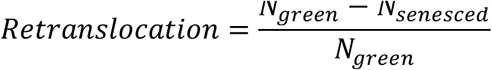

Bark thickness was calculated as a scaling coefficient relative to stem diameter (*β*)

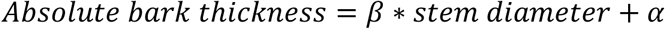

To test the potential for traits to predict the response of trees to fire, we fit linear mixed-effects models with the CWM modifying fire effects but allowing for main effects of fire. For example,

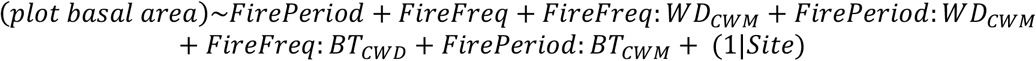

To test how fire influenced the trait composition of the community we fit mixed-effects models to test the effect of both fire as well as environmental factors in explaining the community weighted mean trait values.

We do not include an independent effect of either wood density or bark thickness because we are primarily concerned with how they may modify fire effects.

We also tested for whether the symbiotic strategies differed in their traits. To do so, we assigned symbiotic strategies and the dominant ecosystem in which they occurred to different species. We then analyzed linear models incorporating symbiotic strategy and ecosystem type as additive effects.

Table S1: List of sites with key meta-data. Cont=continent (AU=Australia, NA=North America, SA=South America, AF=Africa). Vegetation type present in broad categories (NL=needleleaf, BL=broadleaf) and the families of the dominant tree species. Sites with a pine-dominated ecosystem that can change from pine to oak depending on fire regime are noted. Number of plots is the total within the entire site. Duration is the number of years over which fire frequencies have differed across plots. Frequency is in # fires yr^-1^. Prior conditions describe the ecosystem type at the beginning of the ecosystem, whether the site experienced regular burning prior to the experiment and if not, how long it had remained unburned (reintroduction burns).

*In attached document*.

**Table S2:** Meta-analysis statistics. The sample size indicates true replicates. The top section analyzes basal area, the bottom analyzes stem abundance.

| **Variable**<br>**measured** | **Fire**<br>**comparison** | *Response*<br>*ratio* | *Standard*<br>*error* | *Z value* | *p value* | *Lower CI* | *Upper CI* |
| --- | --- | --- | --- | --- | --- | --- | --- |
| Basal area | Low vs. high |  |  |  |  |  |  |
|  | n=22 | -0.78 | 0.22 | -3.53 | 0.0004 | -1.22 | -0.35 |
|  | Low vs. mid |  |  |  |  |  |  |
|  | n=16 | -0.40 | 0.12 | -3.41 | 0.0006 | -0.63 | -0.17 |
|  | Mid vs. high |  |  |  |  |  |  |
|  | n=16 | -0.43 | 0.23 | -1.86 | 0.0632 | -0.88 | 0.02 |
| Stem<br>abundance | Low vs. high |  |  |  |  |  |  |
|  | n=23 | -0.58 | 0.19 | -3.13 | 0.002 | -0.94 | -0.22 |
|  | Low vs. mid |  |  |  |  |  |  |
|  | n=17 | -0.25 | 0.12 | -2.12 | 0.034 | -0.48 | -0.02 |
|  | Mid vs. high |  |  |  |  |  |  |
|  | n=18 | -0.33 | 0.20 | -1.65 | 0.0985 | -0.73 | 0.06 |

**Table S3:**
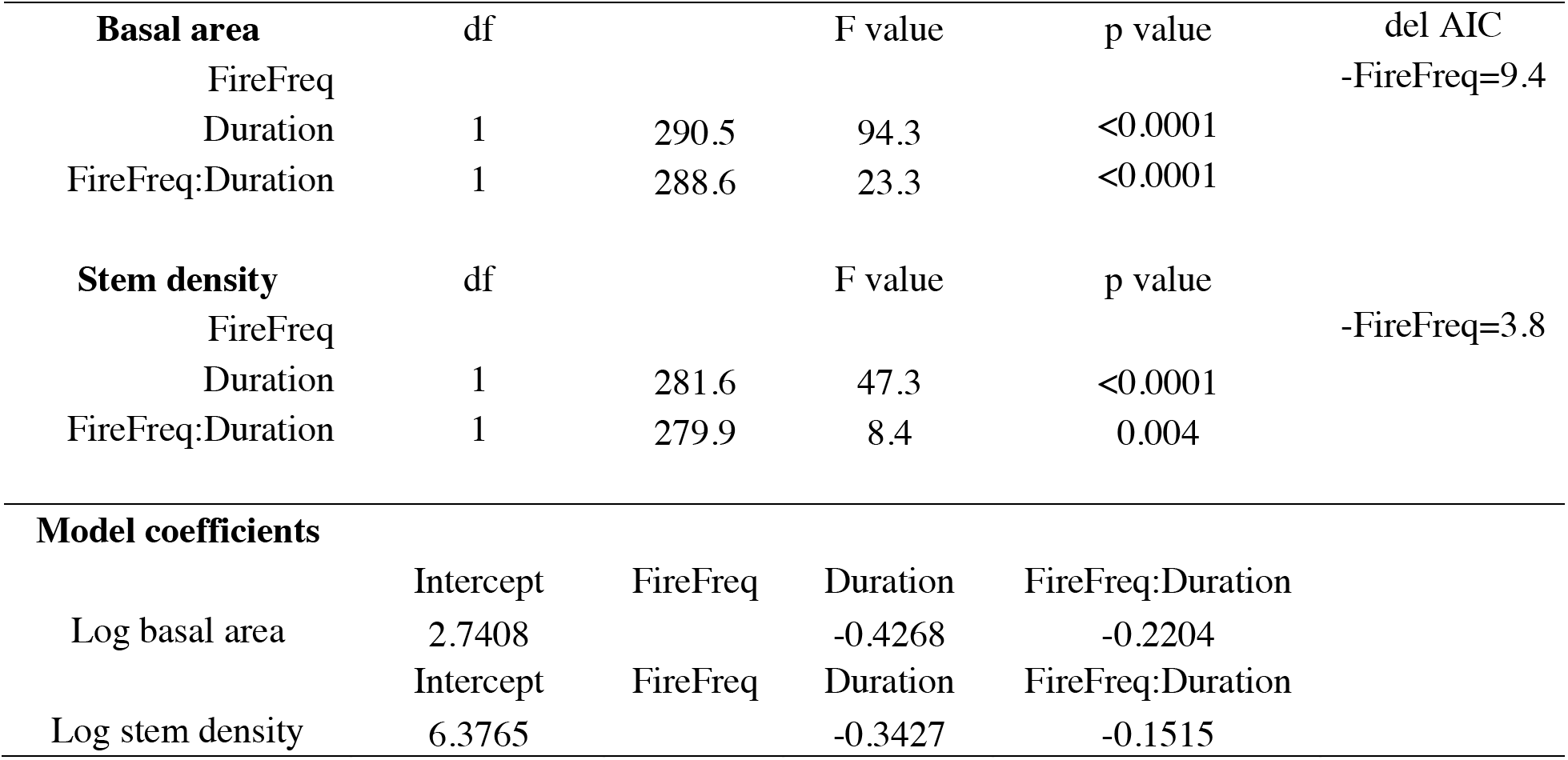
Results from mixed-effects model fit to log basal area and stem density (ANOVA for significance of terms, and then fitted model coefficients) testing the effect of fire frequency (FireFreq), the length of time plots were exposed to different frequencies (Duration) and their interaction (FireFreq:Duration). The means and standard deviations used to re-scale the data were: Basal area: fire frequency, mean=0.34, standard deviation=0.32; duration of experiment, mean=28, standard deviation=19. Stem density: fire frequency, mean=0.35, standard deviation=0.33; duration of experiment, mean=29, standard deviation=19. Units for frequency are fires per-year and duration are years. The main effect of fire frequency was dropped from the top model based on the AIC being lower.

**Table S4:** Results from mixed-effects model fit to log basal area a) ANOVA for significance of terms, b) fitted model coefficients, and c) change in the model AIC with altered additions and removals. All analyses performed on mean centered and standard deviation scaled data for continuous variables with site as a random intercept. ANOVA uses Satterthwaite’s method to estimate degrees of freedom. Colon denotes interactions. Variable abbreviations are: FireFreq= fire frequency (fires yr^-1^), Veg=vegetation type (needleleaf forest, broadleaf forest, savanna), MAT=mean annual temperature (°C), PWQ=precipitation in wet quarter (mm), PDQ=precipitation in dry quarter (mm), Duration=length of time plots have experienced the repeated burning regime (years). For the fitted model coefficients, the intercept gives the value for broadleaf forest (so to calculate the basal area in a savanna, you would exponentiate the sum of the coefficient of “VegSavanna” and the intercept). See Figures 3, S7 for the effects. Independent effects of PWQ, PDQ, and Continent were not included in the model because the models did not pass the criterion that an improved model needed to have a >2 AIC difference. C) Sensitivity of model to changes in terms illustrates what happens when the model only includes interactions and the effect of adding or removing independent effects, as well as the interactions between fire and soil.

| a) Type III ANOVA |  | df | F value | P value | Mean | SD |
| --- | --- | --- | --- | --- | --- | --- |
| FireFreq | 1 | 287.5 | 29.0 | <0.001 | 0.34 | 0.32 |
| Veg | 2 | 17.8 | 1.9 | 0.172 |  |  |
| MAT | 1 | 18.6 | 2.7 | 0.115 |  |  |
| Continent | 3 | 19.2 | 4.2 | 0.020 | 14 | 4 |
| FireFreq:Veg | 2 | 279.7 | 14.0 | <0.001 |  |  |
| FireFreq:PWQ | 1 | 285.1 | 50.3 | <0.001 | 375 | 149 |
| FireFreq:PDQ | 1 | 285.6 | 7.5 | 0.007 | 133 | 108 |
| FireFreq:MAT | 1 | 280.8 | 16.8 | <0.001 | 14 | 4 |
| FirePeriod:Continent | 4 | 283.4 | 7.2 | <0.001 | 28 | 19 |

| b) Fitted model coefficients |  |  |  |  |  |
| --- | --- | --- | --- | --- | --- |
| Variable | Estimate | SE | df | t value | P value |
| (Intercept) | 1.8 | 0.6 | 17.2 | 2.9 | 0.011 |
| FireFreq | -0.1 | 0.1 | 284.7 | -1.5 | 0.135 |
| VegetationNeedleleaf | 0.2 | 0.3 | 18.5 | 0.8 | 0.451 |
| VegetationSavgrass | -0.9 | 0.6 | 17.1 | -1.5 | 0.142 |
| MAT | -0.2 | 0.1 | 18.6 | -1.7 | 0.115 |
| ContinentAustralia | 1.6 | 0.7 | 17.2 | 2.4 | 0.025 |
| ContinentNorthAmer | 1.1 | 0.6 | 17.1 | 1.9 | 0.078 |
| ContinentSouthAmer | 1.7 | 0.6 | 23.2 | 2.8 | 0.010 |
| FireFreq:VegetationNeedleleaf | 0.1 | 0.1 | 278.8 | 1.0 | 0.310 |
| FireFreq:VegetationSavgrass | -0.5 | 0.1 | 281.1 | -3.5 | <0.001 |
| FireFreq:PWQ | -0.4 | 0.1 | 285.1 | -7.1 | <0.001 |
| FireFreq:PDQ | 0.2 | 0.1 | 285.6 | 2.7 | 0.007 |
| FireFreq:MAT | 0.2 | 0.0 | 280.8 | 4.1 | 0.000 |
| FirePeriod:ContinentAfrica | -0.2 | 0.1 | 285.1 | -2.1 | 0.039 |
| FirePeriod:ContinentAustralia | 0.3 | 0.1 | 283.2 | 1.9 | 0.059 |
| FirePeriod:ContinentNorthAmer | -0.2 | 0.1 | 291.3 | -4.3 | <0.001 |
| FirePeriod:ContinentSouthAmer | -0.2 | 0.2 | 274.7 | -0.8 | 0.418 |

*\*\*Intercept using broadleaf for vegetation*

c) Sensitivity of model terms
| *Model* | *df* | *AIC* | *ΔAIC* |
| --- | --- | --- | --- |
| ~FireFreq+Veg+MAT+Veg:FireFreq+PWQ:FireFreq+PDQ:FireFreq |  |  | *Top* |
| +MAT:FireFreq+Continent:Duration+Continent | 19 | 556.3 | *model* |
| Only interactions | 13 | 582.0 | 25.76 |
| +PWQ | 17 | 561.0 | 4.70 |
| +PDQ | 17 | 566.4 | 10.18 |
| -MAT | 16 | 567.3 | 11.03 |
| -VegType | 15 | 589.1 | 32.83 |
| -Continent | 16 | 562.8 | 6.5394 |
| +SoilType | 26 | 559.3 | 3.0709 |
| +%C | 20 | 561.4 | 5.1847 |
| +SoilType:FireFreq | 23 | 581.3 | 25.06 |
| +%C:FireFreq | 17 | 569.1 | 25.76 |

**Table S5:**
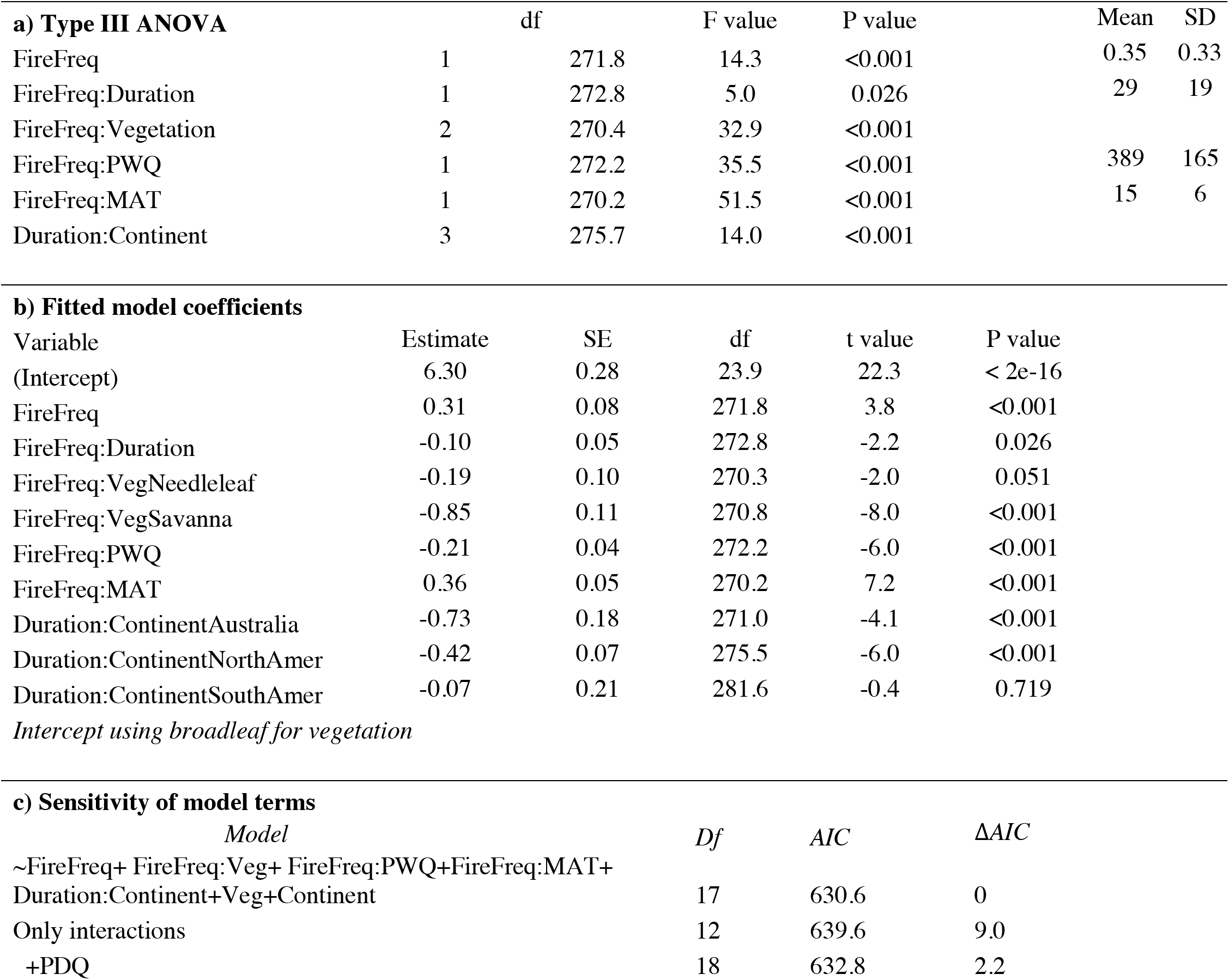
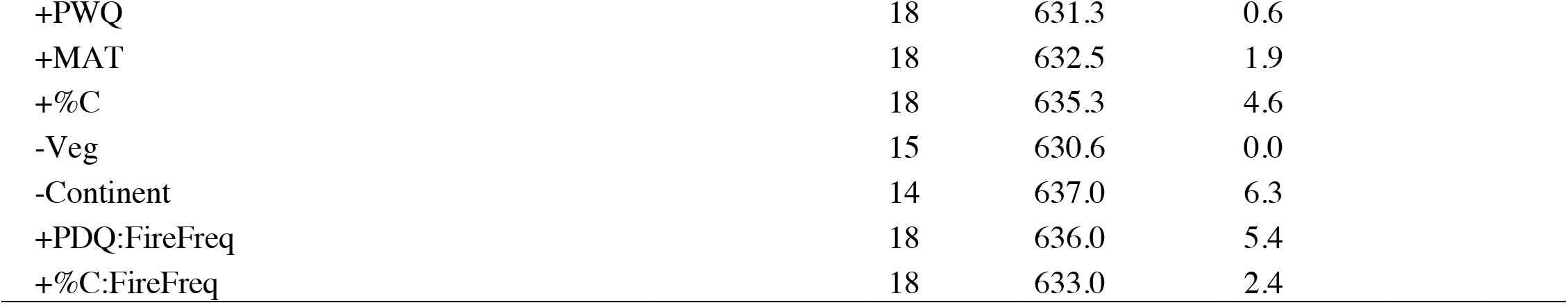
Results from mixed-effects model fit to log stem density a) ANOVA for significance of terms, b) fitted model coefficients, and c) sensitivity of model terms. All analyses performed on mean centered and standard deviation scaled data for continuous variables with site as a random intercept. ANOVA uses Satterthwaite’s method to estimate degrees of freedom. Colons denote interactions. Variable abbreviations are: FireFreq= fire frequency (fires yr^-1^), Veg=vegetation type (needleleaf forest, broadleaf forest, savanna), MAT=mean annual temperature (°C), PWQ=precipitation in wet quarter (mm), Duration=length of time plots have experienced the repeated burning regime (years). For the fitted model coefficients, the intercept gives the value for broadleaf forest (so to calculate the basal area in a savanna, exponentiate the sum of the coefficient of “VegSavanna” and the intercept). See Figures S7, S9 for effects. Model terms are presented relative to the top model (given in the first row), the only interactions refers to all main effects removed.

| **a) Type III ANOVA** |  | df | F value | P value | Mean | SD |
| --- | --- | --- | --- | --- | --- | --- |
| FireFreq | 1 | 271.8 | 14.3 | <0.001 | 0.35 | 0.33 |
| FireFreq:Duration | 1 | 272.8 | 5.0 | 0.026 | 29 | 19 |
| FireFreq:Vegetation | 2 | 270.4 | 32.9 | <0.001 |  |  |
| FireFreq:PWQ | 1 | 272.2 | 35.5 | <0.001 | 389 | 165 |
| FireFreq:MAT | 1 | 270.2 | 51.5 | <0.001 | 15 | 6 |
| Duration:Continent | 3 | 275.7 | 14.0 | <0.001 |  |  |
| **b) Fitted model coefficients** |  |  |  |  |  |  |
| Variable | Estimate | SE | df | t value | P value |  |
| (Intercept) | 6.30 | 0.28 | 23.9 | 22.3 | < 2e-16 |  |
| FireFreq | 0.31 | 0.08 | 271.8 | 3.8 | <0.001 |  |
| FireFreq:Duration | -0.10 | 0.05 | 272.8 | -2.2 | 0.026 |  |
| FireFreq:VegNeedleleaf | -0.19 | 0.10 | 270.3 | -2.0 | 0.051 |  |
| FireFreq:VegSavanna | -0.85 | 0.11 | 270.8 | -8.0 | <0.001 |  |
| FireFreq:PWQ | -0.21 | 0.04 | 272.2 | -6.0 | <0.001 |  |
| FireFreq:MAT | 0.36 | 0.05 | 270.2 | 7.2 | <0.001 |  |
| Duration:ContinentAustralia | -0.73 | 0.18 | 271.0 | -4.1 | <0.001 |  |
| Duration:ContinentNorthAmer | -0.42 | 0.07 | 275.5 | -6.0 | <0.001 |  |
| Duration:ContinentSouthAmer | -0.07 | 0.21 | 281.6 | -0.4 | 0.719 |  |
| *Intercept using broadleaf for vegetation* |  |  |  |  |  |  |
| **c) Sensitivity of model terms** |  |  |  |  |  |  |
| *Model* |  | *Df* | *AIC* | *ΔAIC* |  |  |
| ~FireFreq+ FireFreq:Veg+ FireFreq:PWQ+FireFreq:MAT+ Duration:Continent+Veg+Continent |  | 17 | 630.6 | 0 |  |  |
| Only interactions |  | 12 | 639.6 | 9.0 |  |  |
| +PDQ |  | 18 | 632.8 | 2.2 |  |  |

|  |  |  |  |
| --- | --- | --- | --- |
| +PWQ | 18 | 631.3 | 0.6 |
| +MAT | 18 | 632.5 | 1.9 |
| +%C | 18 | 635.3 | 4.6 |
| -Veg | 15 | 630.6 | 0.0 |
| -Continent | 14 | 637.0 | 6.3 |
| +PDQ:FireFreq | 18 | 636.0 | 5.4 |
| +%C:FireFreq | 18 | 633.0 | 2.4 |

**Table S6:** Results from mixed-effects models testing how: a) wood traits modified fire effects on tree basal area and b) fire altered the trait values within plots. Whether the site was a savanna, broadleaf forest, or needleleaf forest was included in the model because of the large difference in wood traits between needleleaf forests and the other ecosystems.

| <u>a) Basal area ~ wood traits</u> | df |  | F value | P value |
| --- | --- | --- | --- | --- |
| FireFreq | 1 | 154.4 | 12.9 | <0.001 |
| FireFreq:Bark | 1 | 154.3 | 5.7 | 0.018 |
| FireFreq:WD | 1 | 154.1 | 12.9 | <0.001 |
| <u>b) Wood traits ~ fire</u> |  |  |  |  |
| *Bark* |  |  |  |  |
| Duration | 1 | 157.3 | 0.4 | 0.520 |
| FireFreq | 1 | 156.3 | 0.2 | 0.624 |
| Vegetation | 2 | 12.9 | 7.9 | 0.006 |
| *Wood density* |  |  |  |  |
| Duration | 1 | 156.4 | 5.4 | 0.022 |
| Vegetation | 2 | 13.0 | 9.8 | 0.003 |

**Table S7:** Results from mixed-effects models testing the effect of fire on nitrogen (N) and phosphorus (P) concentrations in green and senesced leaves, the proportion of N and P retranslocated before senescence, and root N and P concentrations. Vegetation type was included as a term given the strong differences in traits between needleleaf vs. broadleaf trees.

| Green N | df |  | F value | P value | Green P | df |  | F value | P value |
| --- | --- | --- | --- | --- | --- | --- | --- | --- | --- |
| Duration | 1 | 158.9 | 13.8 | <0.001 | Duration | 1 | 157.3 | 11.8 | <0.001 |
| Vegetation | 2 | 12.7 | 33.6 | <0.001 | Vegetation | 2 | 12.6 | 12.4 | 0.001 |
| Senesced N |  |  |  |  | Senesced P |  |  |  |  |
| Duration | 1 | 158.9 | 19.7 | <0.001 | Duration | 1 | 158.3 | 21.3 | <0.001 |
| Vegetation | 2 | 12.4 | 19.1 | <0.001 | Vegetation | 2 | 12.5 | 27.2 | <0.001 |
| Retrans N |  |  |  |  | Retrans P |  |  |  |  |
| Duration | 1 | 157.8 | 14.2 | <0.001 | Duration | 1 | 159.8 | 24.1 | <0.001 |
| Vegetation | 2 | 12.3 | 3.0 | 0.088 | Vegetation | 2 | 12.5 | 43.3 | <0.001 |
| Root N |  |  |  |  | Root P |  |  |  |  |
| Duration | 1 | 160.0 | 14.3 | <0.001 | Duration | 1 | 157.2 | 3.1 | 0.082 |
| Vegetation | 2 | 12.6 | 30.7 | <0.001 | Vegetation | 2 | 12.4 | 1.1 | 0.378 |

**Table S8:** Results from top mixed-effects model on the relative abundance of trees summed within a symbiotic strategy within a plot with site as a random intercept conducted in North America where taxonomic resolution was the highest (relative basal area was arcsine transformed). The statistics were only run on ectomycorrhizal and arbuscular mycorrhizal groups because they were sufficiently abundant across plots, but all other strategies (ericoid, non-mycorrhizal, nitrogen fixer) were included in relative basal area calculation.

| Variable | df | df | F value | P value |
| --- | --- | --- | --- | --- |
| Symb | 1 | 138.3 | 49.1 | <0.001 |
| Symb:Soil C | 2 | 34.4 | 23.4 | <0.001 |
| Symb:FireFreq | 2 | 140.1 | 7.8 | <0.001 |
| Symb:MAT | 2 | 18.6 | 12.5 | <0.001 |
| Symb:Family | 4 | 14.0 | 4.4 | 0.017 |
| Symb:Duration:FireFreq | 2 | 135.0 | 4.0 | 0.021 |

**Table S9:** Type III ANOVAs on linear models testing the differences in tissue stoichiometry between the symbiotic strategy grouped in the different ecosystem vegetation types (broadleaf forest, needleleaf forest, or savanna). Inferred statistics are using phylogenetic relationships to infer trait values for species with missing data (see supporting information and^46^) while observed are based on direct trait measurements. The inferred vs. observed do not refer to the classification of mycorrhizal type.

| **Inferred** |  |  |  | **Observed** |  |  |  |
| --- | --- | --- | --- | --- | --- | --- | --- |
| N green |  |  |  | N green |  |  |  |
| n=61 | DF | F value | p value | n=48 | Df | F value | p value |
| Symb | 1 | 12.4 | 0.001 | Symb | 1 | 4.2 | 0.046 |
| Veg | 2 | 5.4 | 0.007 | Veg | 2 | 4.0 | 0.025 |
| N senesced |  |  |  | N senesced |  |  |  |
| n=61 | Df | F value | p value | n=25 | Df | F value | p value |
| Symb | 1 | 90.8 | 0.000 | Symb | 1 | 1.1 | 0.310 |
| Veg | 2 | 4.5 | 0.016 | Veg | 2 | 1.5 | 0.237 |
| N roots |  |  |  | N roots |  |  |  |
| n=61 | Df | F value | p value | n=31 | Df | F value | p value |
| Symb | 1 | 15.8 | 0.000 | Symb | 1 | 0.6 | 0.427 |
| Veg | 2 | 1.8 | 0.181 | Veg | 2 | 0.7 | 0.489 |
| P green |  |  |  | P green |  |  |  |
| n=61 | DF | F value | p value | n=38 | Df | F value | p value |
| Symb | 1 | 8.5 | 0.005 | Symb | 1 | 3.9 | 0.054 |
| Veg | 2 | 1.8 | 0.176 | Veg | 2 | 5.7 | 0.007 |
| P senesced |  |  |  | P senesced |  |  |  |
| n=61 | DF | F value | p value | n=21 | Df | F value | p value |
| Symb | 1 | 27.8 | 0.000 | Symb | 1 | 6.6 | 0.018 |
| Veg | 2 | 6.1 | 0.004 | Veg | 2 | 1.6 | 0.216 |
| P roots |  |  |  | P roots |  |  |  |
| n=61 | DF | F value | p value | n=9 | Df | F value | p value |
| Symb | 1 | 11.0 | 0.002 | Symb | 1 | 9.3 | 0.014 |
| Veg | 2 | 0.6 | 0.572 | Veg | 2 | 1.2 | 0.346 |

**Table S10:** Pearson correlation coefficients between climate variables., PWQ=precipitation in wet quarter (mm), PDQ=precipitation in dry quarter (mm), MAT=mean annual temperature (°C), MAP=mean annual precipitation (mm yr^-1^). Data derived from WorldClim.

|  | PDQ | PWQ | MAT | MAP | Aridity |
| --- | --- | --- | --- | --- | --- |
| PDQ |  | 0.28 | -0.05 | **0.82** | **0.88** |
| PWQ | 0.28 |  | 0.44 | **0.77** | 0.54 |
| MAT | -0.05 | 0.44 |  | 0.19 | -0.04 |
| MAP | **0.82** | **0.77** | 0.19 |  | **0.91** |
| Aridity | **0.88** | 0.54 | -0.04 | **0.91** |  |

### Supplemental figures

**Figure S1:**
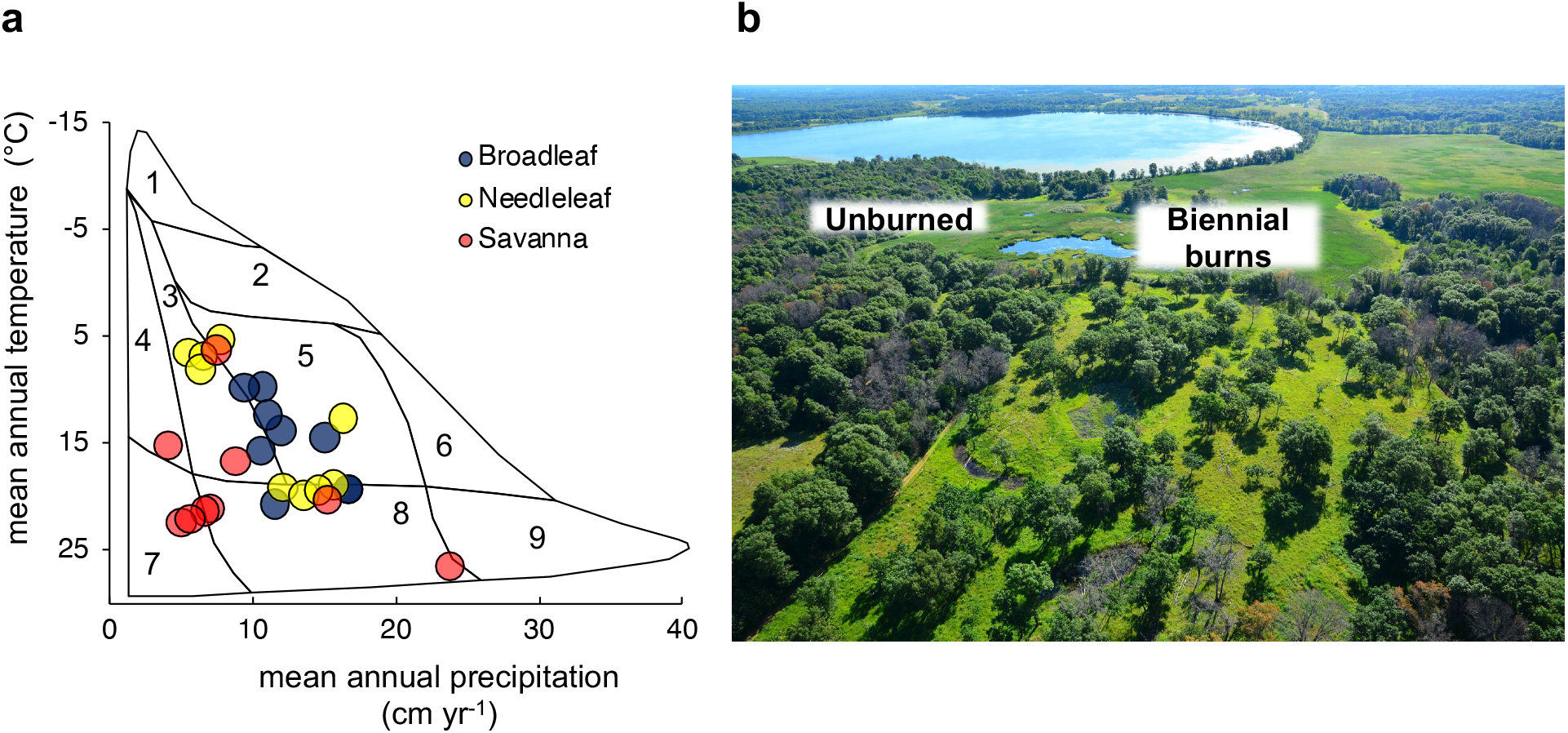
a) distribution of sites in climate space overlying Whittaker’s biome distribution ^81^. (1=tundra, 2=boreal forest, 3=woodland/shrubland, 4=temperate grassland/desert, 5=temperate forest, 6=temperate rainforest, 7=subtropical desert, 8=tropical forest and savanna, 9=tropical rainforest). Dots colored according to broad vegetation type category. Plots span a mean annual temperate range from 5.2-27.3° C and a mean annual precipitation range from 408-2378 mm yr^-1^. b) aerial picture of two different fire treatment plots from Cedar Creek, a temperate oak savanna, where different fire frequencies have created a stark biome boundary between forests in unburned plots and savannas in biennial burn plots (Pellegrini et al. 2019).

**Figure S2:**
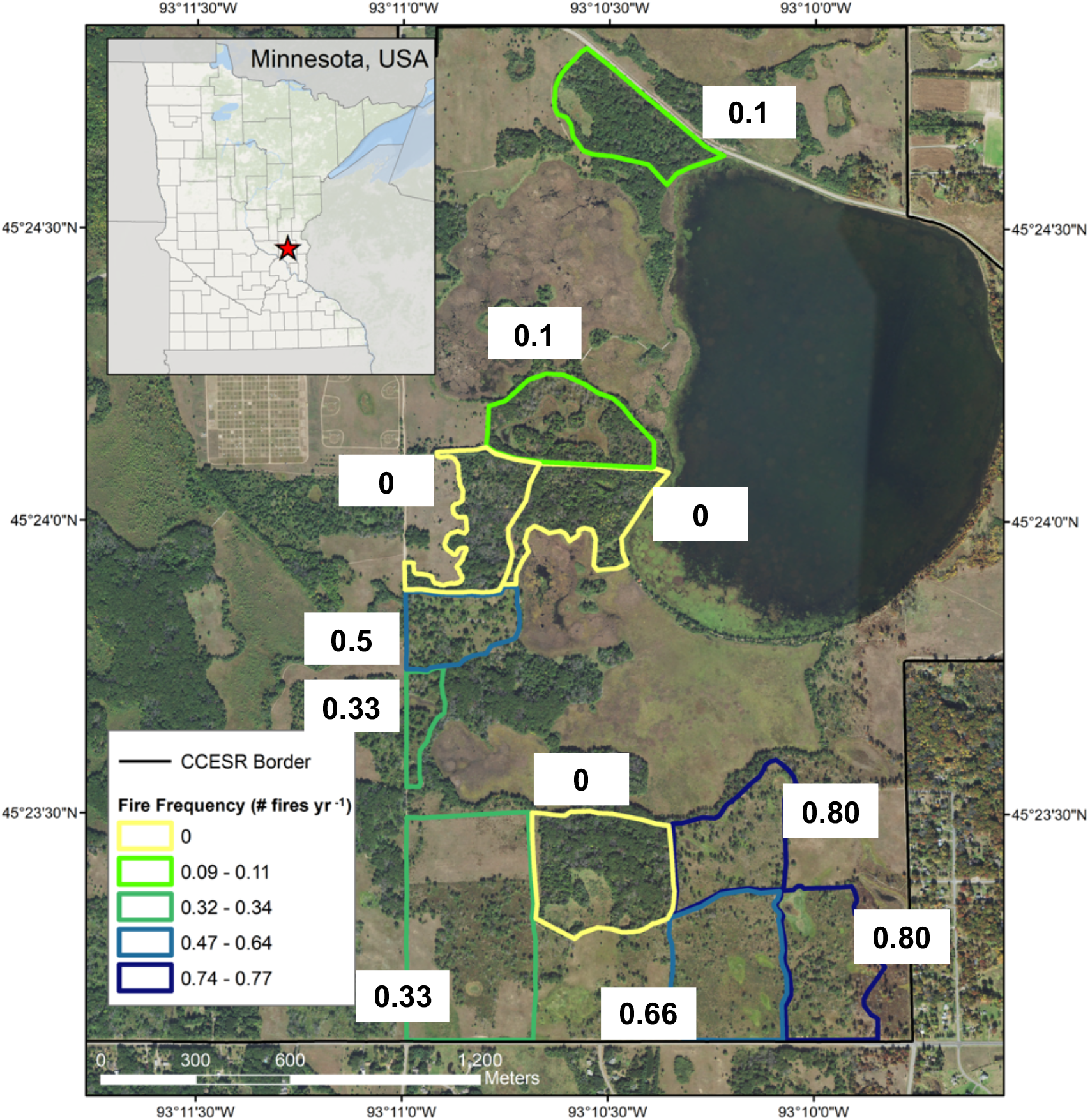
Example of the experimental layout of a fire manipulation experiment taken from Cedar Creek (a temperate savanna in Minnesota, USA), where fires have been manipulated since 1964. Aerial imagery (taken in 2017) from the National Agriculture Imagery Program from the Farm Service Agency. Plots are outlined with a color corresponding to their fire frequencies expressed in terms of number of fires per year (e.g. 0.33 is one fire every 3 years).

**Figure S3:**
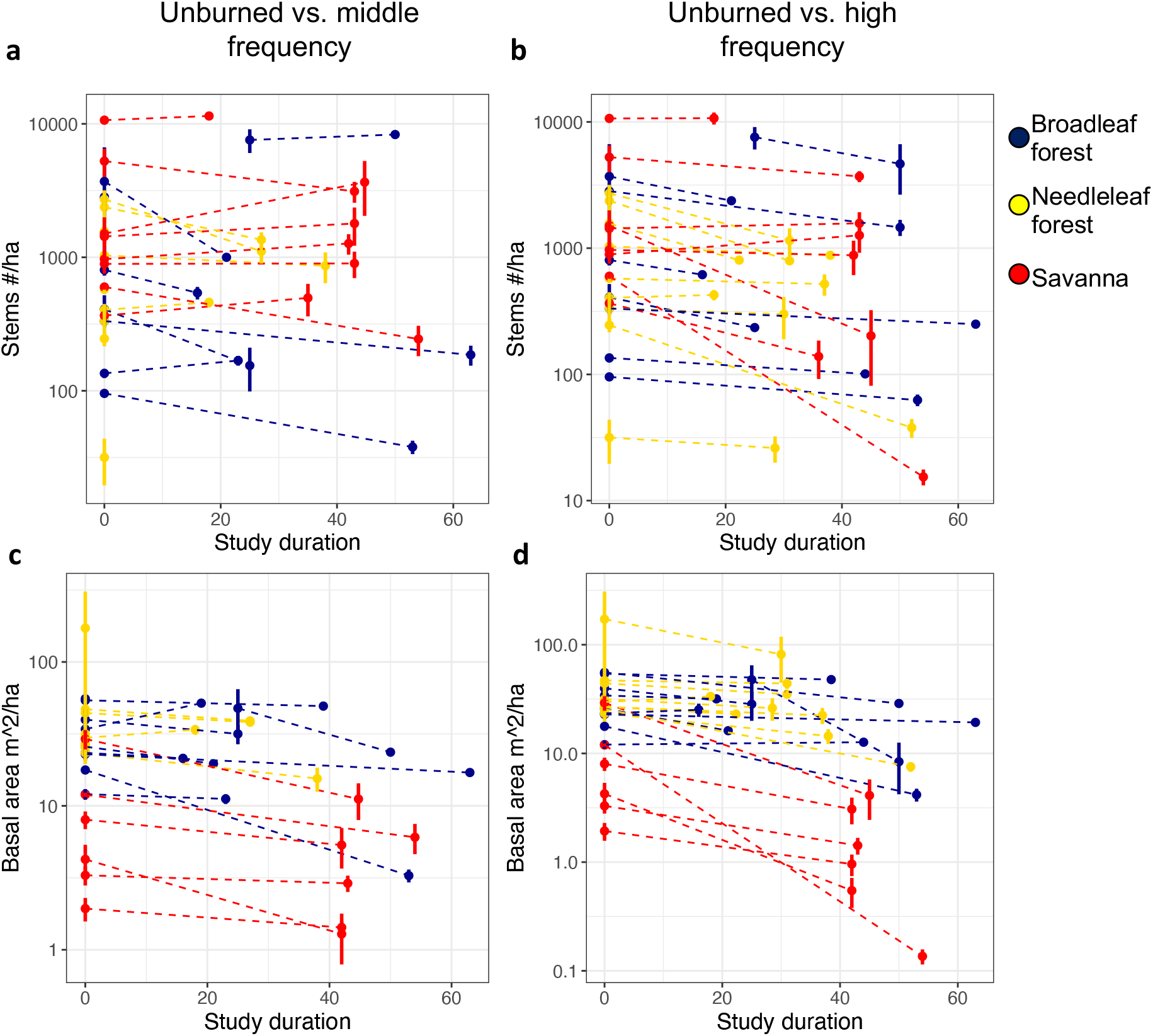
Untransformed data on stem density (**a-b**) and basal area (**c-d**) as a function of the duration that plots have been exposed to burning in the experiment (0=unburned plots). Each dot represents a site and the dashed lines connect treatments within sites. Columns represent two sets of fire frequency contrasts comparing unburned vs. the intermediate frequency in **a** and **c**, and unburned vs. the high frequency in **b** and **d** (levels defined based on treatments within sites). Dots and bars based on mean and standard error calculated across the replicate plots within a fire treatment in a site. Note y-axis is on a log10 scale.

**Figure S4:**
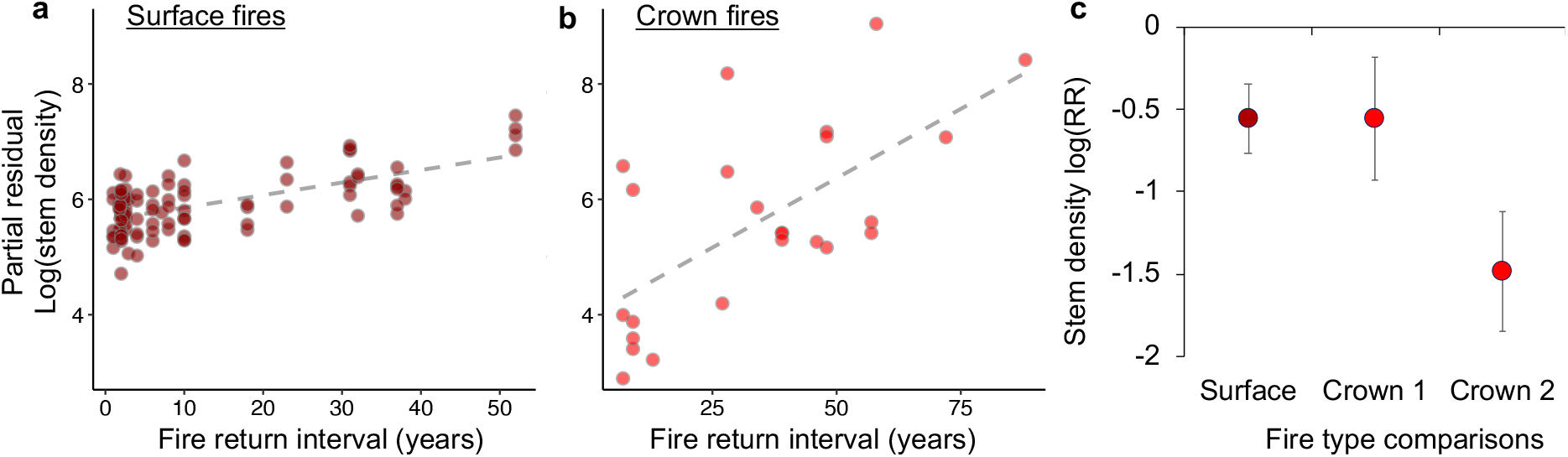
Comparison between fire types (surface in **a**, F_1,94.3_=50.6, p<0.001, and crown in **b**, F_1,21_=10.3, p=0.004) in needleleaf forests with fire expressed in terms of return period (crown fire plots are all 12 years postfire, data subset to include short-interval burn plots). **c**) illustrates the mean response ratios +/- standard error for the fire types with crown fires split into high (>2,400 m) and low (<2,400) elevation sites (Crown 1 and Crown 2, respectively). Analyses were robust to considering surface fires in only Western US needleleaf forests: F_1,47.1_=17.2, p=0.001. Response ratios were split into long and short fire return interval plots (Crown 1 and 2, respectively), with the justification for definition of interval in ^23^.

**Figure S5:**
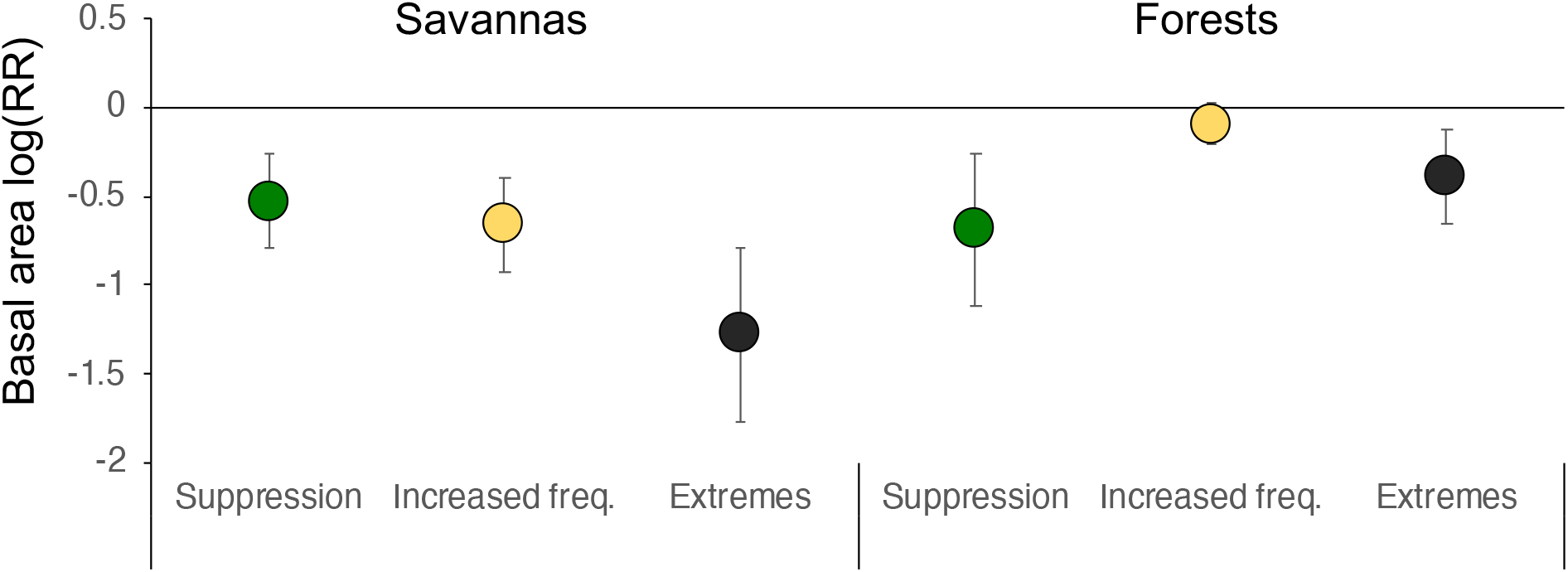
Log response ratios of basal area under different fire history scenarios. In savannas, suppression compares the unburned vs. intermediate burn in a historically burned environment. The increased frequency compares higher than historical frequency with historical frequency. The extremes compare the highest frequency vs. suppression. The main difference in forests is the increased frequency, which is the reintroduction of fire into a historically fire suppressed forest.

**Figure S6:**
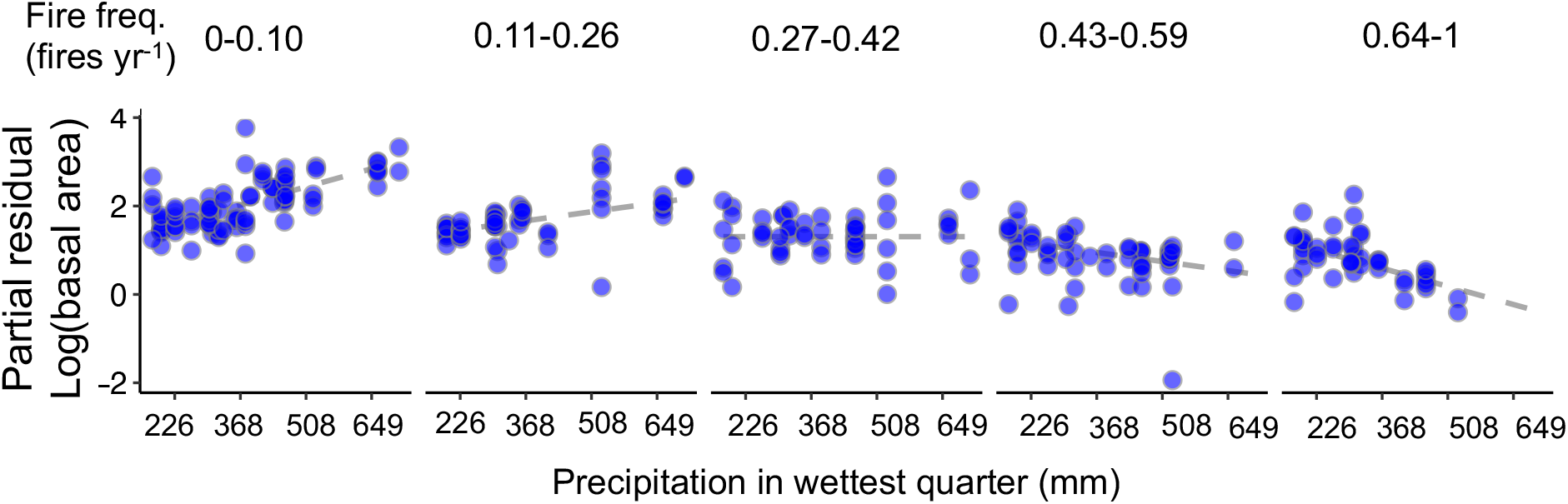
partial residual plot displaying the relationship between loge basal area and precipitation in the wettest quarter cross-sectioned based on fire frequency. This plot is based on the same mixed-effects model presented in Figure 3 and Table S4, just re-arranged to emphasize how precipitation-basal area relationship changes with more frequent burning.

**Figure S7:**
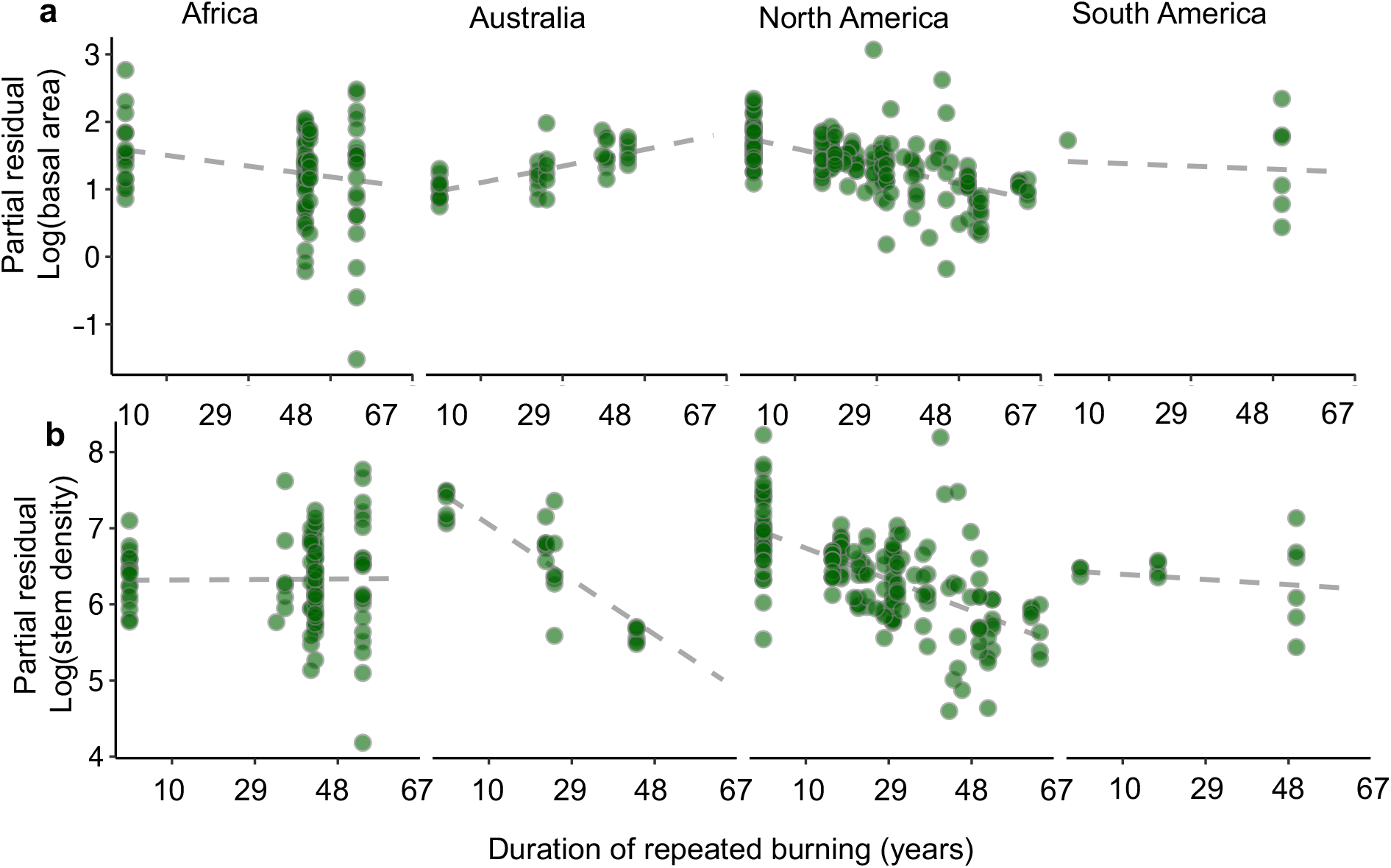
partial residual plot between the length of time plots were exposed to frequent burning and the log basal area (a) and stem density (b) in the different continents (from the main mixed-effects model with site as a random intercept in Tables S4–S5).

**Figure S8:**
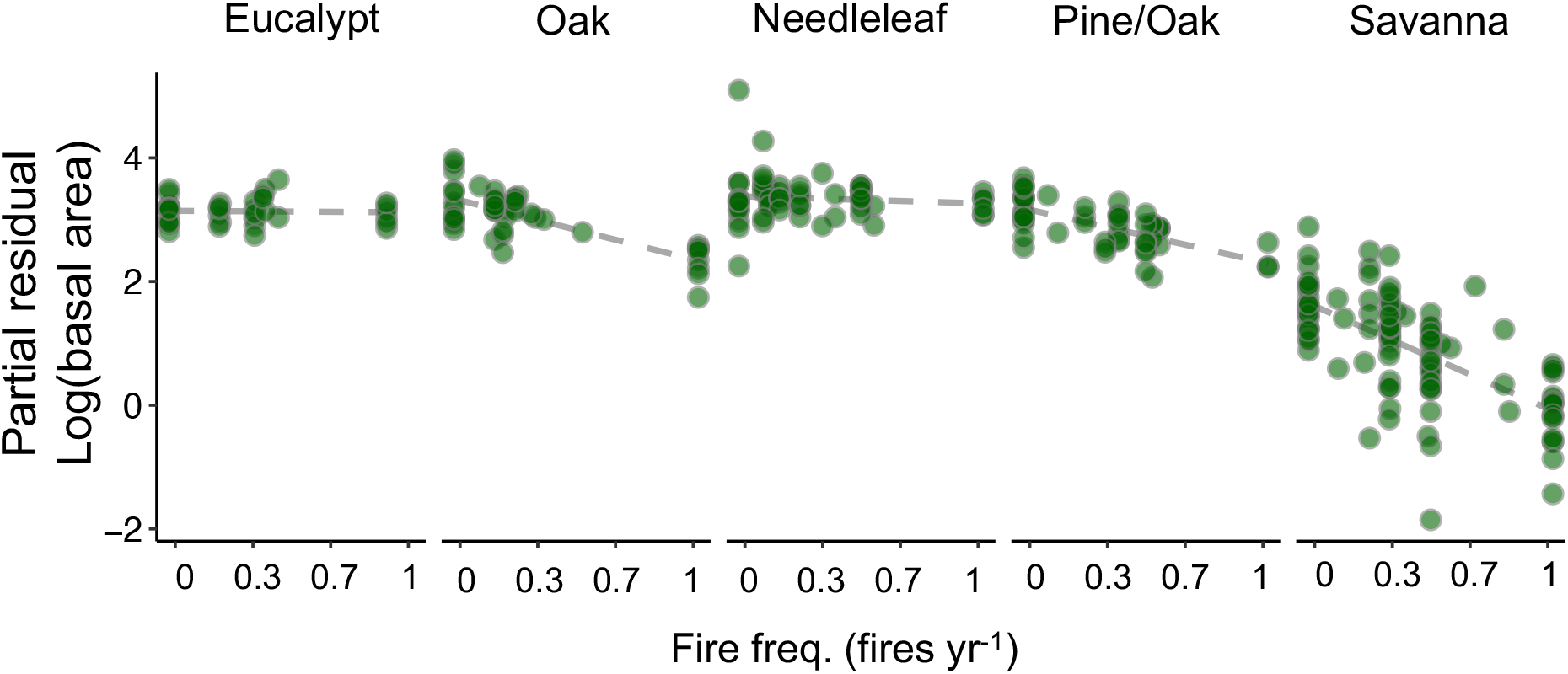
partial residual plot between the length of time plots were exposed to frequent burning and the log basal area in the different sub-vegetation types (from the main mixed-effects model, presented in Table S4 but substituting the broad vegetation effect with the more detailed classification. We found no evidence that accounting for the finer-scale variability in ecosystem classification increased the accuracy of the model or changed our conclusions

**Figure S9:**
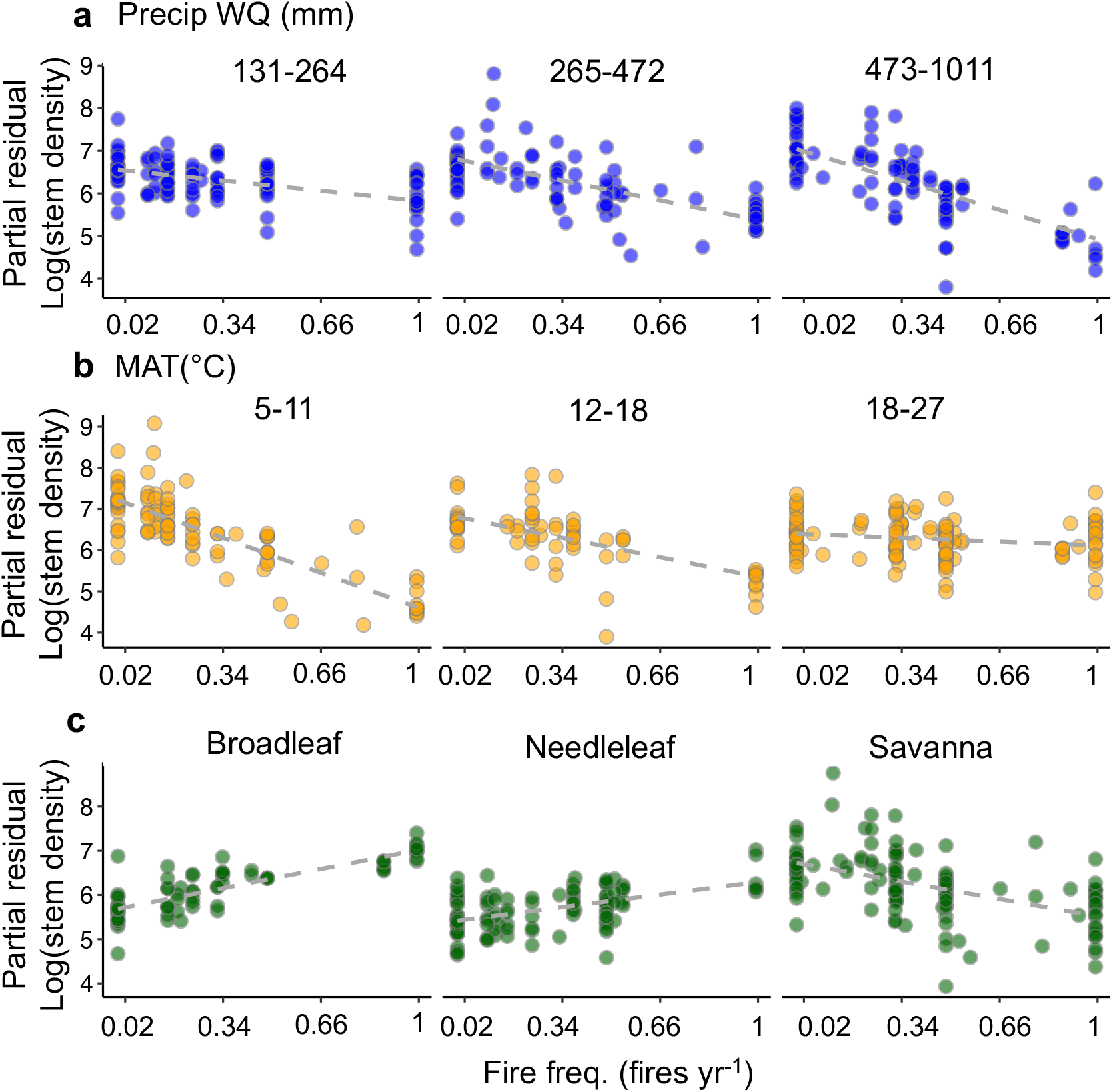
Partial residual plots of the mixed-effects model for stem densities illustrating how fire frequency effects changed according to wet-season precipitation, mean annual temperature, and ecosystem type. Panels structured by standard deviations around the median to visualize the spread (−1, 0,1), PWQ: precipitation in the wet quarter, MAT: mean annual temperature. All model fits are p<0.05 and specific results can be found in Table S5. The predictor variables are mean-centered and standard deviations are scaled to facilitate comparisons of variable influence. In needleleaf and broadleaf forests, stem densities actually increased with more frequent burning initially, but declined with increasing experiment duration, potentially because of increased light availability initially stimulating recruitment of small trees (Figure S7, Table S5). Stem density in African sites changed little through time (Figure S7). The trends in density may reflect the ability of many of the tree species to re-sprout in between fire events^82^.

**Figure S10:**
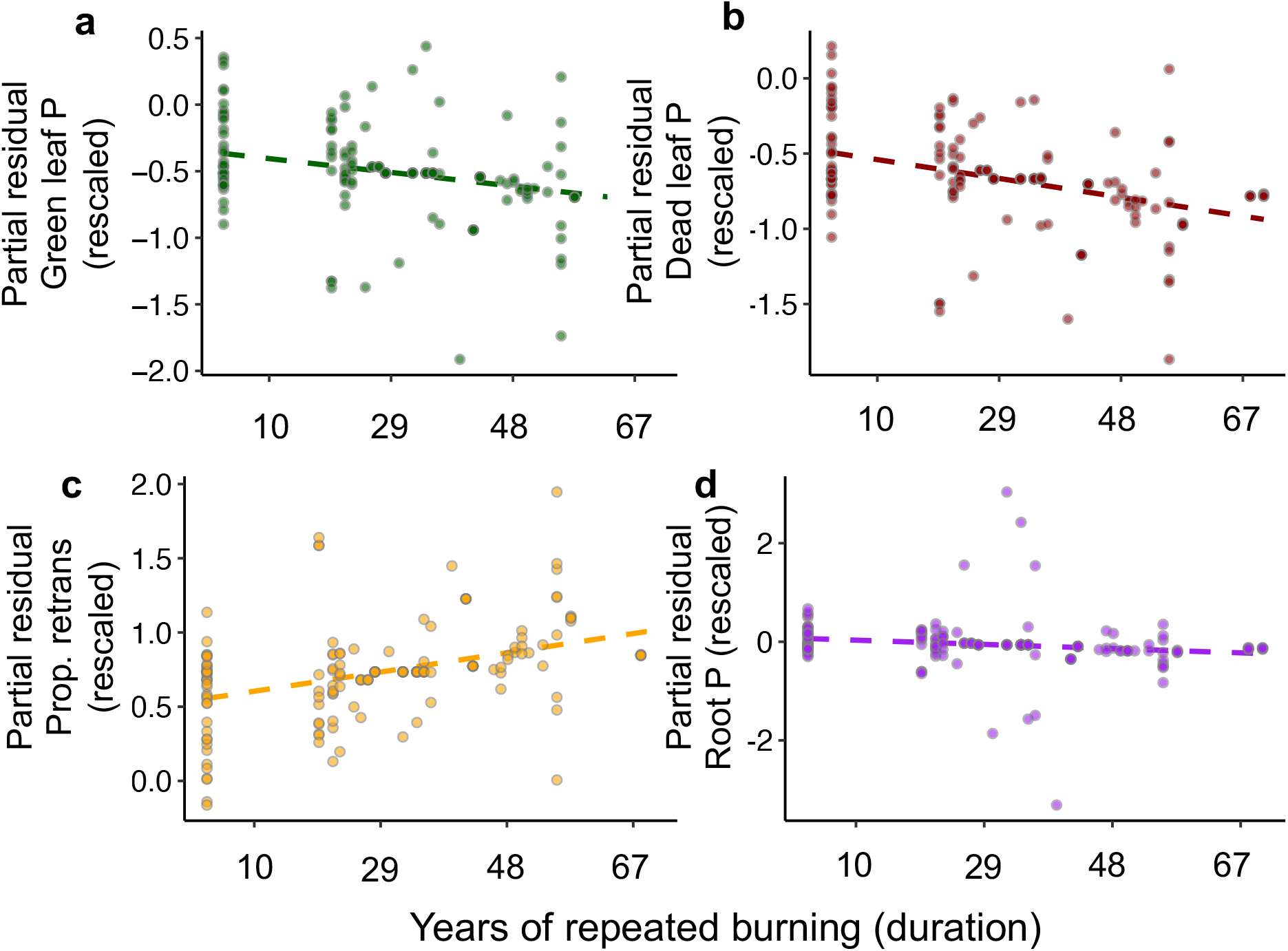
Partial residual plots of the phosphorus (P) stoichiometry of community weighted means as a function of years of repeated burning. Taken from mixed-effects models presented in Table S7. The models include a vegetation type effect. Tissue P is rescaled by subtracting the mean and dividing by the standard deviation.

**Figure S11:**
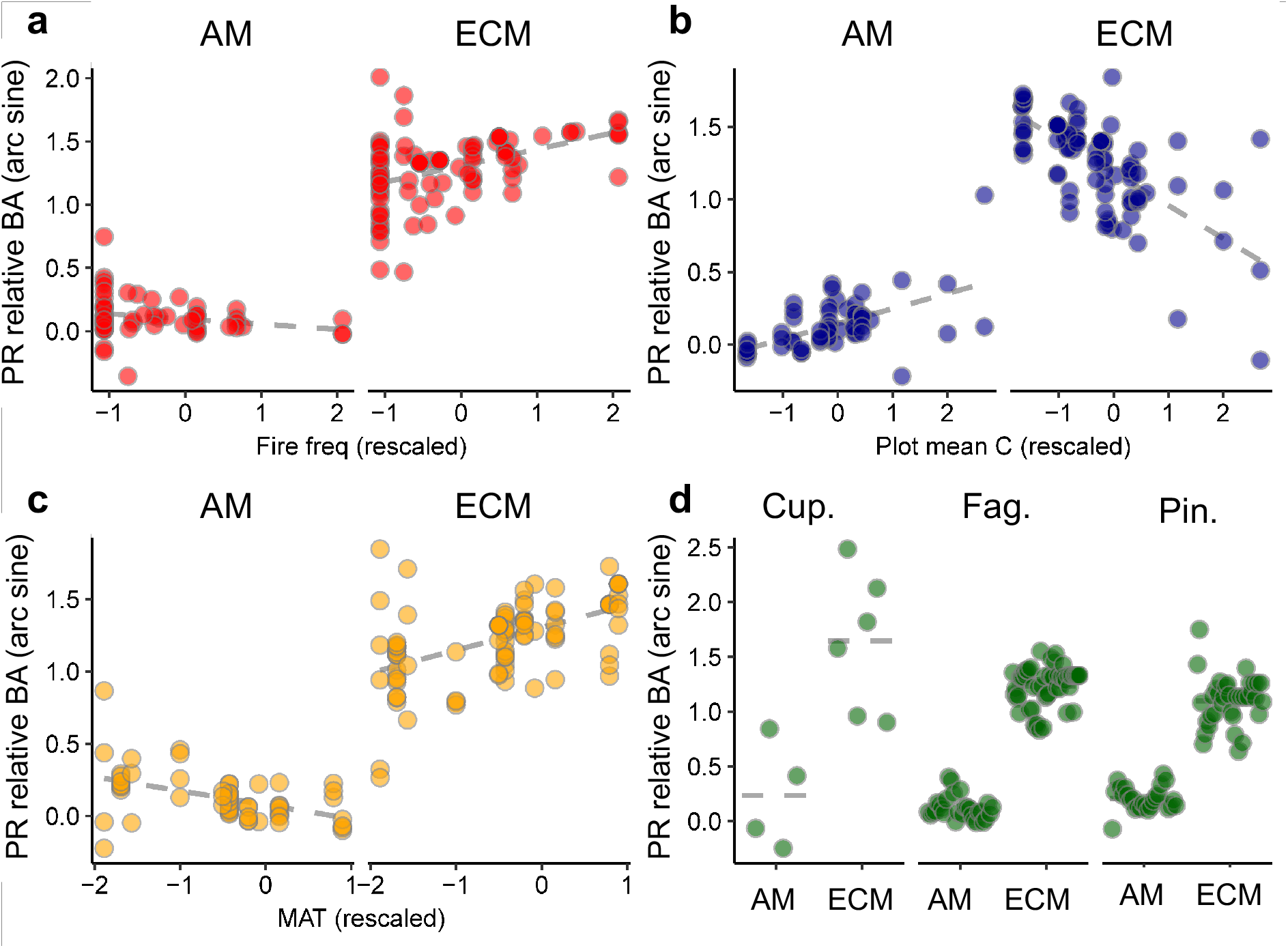
Partial residual (PR) plots of the mixed-effects model between relative basal area different symbiotic groups within a plot (AM=arbuscular mycorrhizal and ECM=ectomycorrhizal on the left- and right-hand side of each panel, respectively). Relative basal area was arcsine transformed. In all panels, the continuous predictor variables were re-scaled by mean centering and dividing by the standard deviation for comparability testing the relationship with fire frequency (a), soil total carbon content (b). mean annual temperature (c). d) illustrates composition across different plant communities grouped based on the family of the dominant tree species (Cup= *Cupressaceae, Fag=Fagaceae, Pin=Pinaceae*). Statistics are given in Table S8.

**Figure S12:**
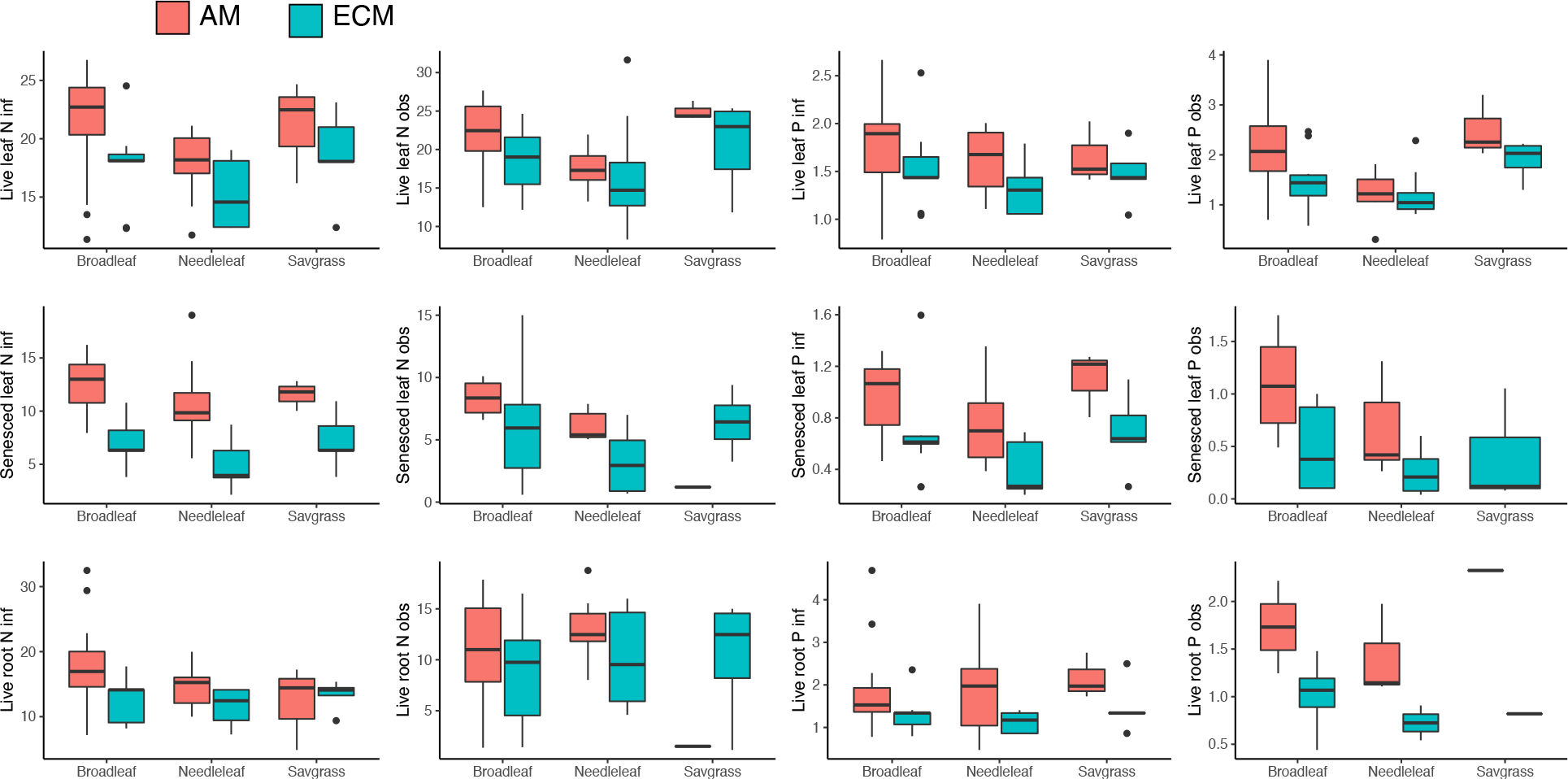
Box and whiskers plot displaying the tissue stoichiometry for tree species averaged within symbiont strategy (AM= arbuscular mycorrhizal; ECM=ectomycorrhizal) and then grouped according to which overall ecosystem type the species generally occurred in (broadleaf forest, needleleaf forest, or a savanna-grassland). For both N and P, we conducted our comparisons using data that were either based on direct observations (“obs” in the y-axis), or inferred via a phylogenetic relatedness statistical filling (“inf” in the y-axis). Statistics are in Table S9.

## References

1. Andela, N. et al. A human-driven decline in global burned area. Science (80-.). 356, 1356–1362 (2017).

2. Westerling, A. L., Hidalgo, H. G., Cayan, D. R. & Swetnam, T. W. Warming and earlier spring increase western US forest wildfire activity. Science (80-.). 313, 940–943 (2006).

3. Turner, M. G. Disturbance and landscape dynamics in a changing world. Ecology 91, 2833–2849 (2010).

4. Higgins, S. I. & Scheiter, S. Atmospheric CO2 forces abrupt vegetation shifts locally, but not globally. Nature 488, 209–212 (2012).

5. Yin, Y. et al. Fire decline in dry tropical ecosystems enhances decadal land carbon sink. Nat. Commun. 11, 1900 (2020).

6. van der Werf, G. R. et al. Global fire emissions estimates during 1997–2016. Earth Syst. Sci. Data 9, 697–720 (2017).

7. Randerson, J. T., Chen, Y., Werf, G. R., Rogers, B. M. & Morton, D. C. Global burned area and biomass burning emissions from small fires. J. Geophys. Res. Biogeosciences 117, (2012).

8. Schoennagel, T. et al. Adapt to more wildfire in western North American forests as climate changes. Proceedings of the National Academy of Sciences vol. 114 4582–4590 (2017).

9. Westerling, A. L., Turner, M. G., Smithwick, E. A. H., Romme, W. H. & Ryan, M. G. Continued warming could transform Greater Yellowstone fire regimes by mid-21st century. Proc. Natl. Acad. Sci. 108, 13165–13170 (2011).

10. Johnstone, J. F. et al. Changing disturbance regimes, ecological memory, and forest resilience. Front. Ecol. Environ. 14, 369–378 (2016).

11. Lewis, T. Very frequent burning encourages tree growth in sub-tropical Australian eucalypt forest. For. Ecol. Manage. 459, 117842 (2020).

12. Peterson, D. W. & Reich, P. B. Prescribed fire in oak savanna: fire frequency effects on stand structure and dynamics. Ecol. Appl. 11, 914–927 (2001).

13. Tilman, D. et al. Fire suppression and ecosystem carbon storage. Ecology 81, 2680–2685 (2000).

14. Pellegrini, A. F. A., Hedin, L. O., Staver, A. C. & Govender, N. Fire alters ecosystem carbon and nutrients but not plant nutrient stoichiometry or composition in tropical savanna. Ecology 96, 1275–1285 (2015).

15. Russell-Smith, J., Whitehead, P. J., Cook, G. D. & Hoare, J. L. Response of Eucalyptus-dominated savanna to frequent fires: lessons from Munmarlary, 1973-1996. Ecol. Monogr. 73, 349–375 (2003).

16. Lehmann, C. E. R. et al. Savanna vegetation-fire-climate relationships differ among continents. Science (80-.). 343, 548–552 (2014).

17. Staver, A. C., Archibald, S. & Levin, S. a. The global extent and determinants of savanna and forest as alternative biome states. Science (80-.). 334, 230–232 (2011).

18. Staver, A. C. Prediction and scale in savanna ecosystems. New Phytol. 219, 52–57 (2018).

19. Higgins, S. I., Bond, J. I. & Trollope, W. S. Fire, resprouting and variability: a recipe for grass-tree coexistence in savanna. J. Ecol. 88, 213–229 (2000).

20. Hoffmann, W. A. et al. Ecological thresholds at the savanna-forest boundary: how plant traits, resources and fire govern the distribution of tropical biomes. Ecol. Lett. 15, 759–768 (2012).

21. Keeley, J. E., Pausas, J. G., Rundel, P. W., Bond, W. J. & Bradstock, R. A. Fire as an evolutionary pressure shaping plant traits. Trends Plant Sci. 16, 406–411 (2011).

22. Brando, P. M. et al. Fire-induced tree mortality in a neotropical forest: the roles of bark traits, tree size, wood density and fire behavior. Glob. Chang. Biol. 18, 630–641 (2012).

23. Schoennagel, T., Turner, M. G. & Romme, W. H. The influence of fire interval and serotiny on postfire lodgepole pine density in Yellowstone National Park. Ecology 84, 2967–2978 (2003).

24. Higgins, S. I. et al. Which traits determine shifts in the abundance of tree species in a fire-prone savanna? J. Ecol. 100, 1400–1410 (2012).

25. Newland, J. A. & DeLuca, T. H. Influence of fire on native nitrogen-fixing plants and soil nitrogen status in ponderosa pine - Douglas-fir forests in western Montana. Can. J. For. Res. 30, 274–282 (2000).

26. Hobbie, S. E. Plant species effects on nutrient cycling: revisiting litter feedbacks. Trends Ecol. Evol. 30, 357–363 (2015).

27. Averill, C. & Hawkes, C. V. Ectomycorrhizal fungi slow soil carbon cycling. Ecol. Lett. 19, 937–947 (2016).

28. Knops, J. M. H., Bradley, K. L. & Wedin, D. A. Mechanisms of plant species impacts on ecosystem nitrogen cycling. Ecol Lett 5, 454–466 (2002).

29. Higgins, S. I. et al. Effects of four decades of fire manipulation on woody vegetation structure in savanna. Ecology 88, 1119–1125 (2007).

30. Pellegrini, A. F. A., Pringle, R. M., Govender, N. & Hedin, L. O. plant biomass and carbon exchange depend on elephant-fire interactions across a productivity gradient in African savanna. J. Ecol. 105, 111–121 (2017).

31. Veenendaal, E. M. et al. On the relationship between fire regime and vegetation structure in the tropics. New Phytol. 218, 153–166 (2018).

32. Guinto, D. F. et al. Soil chemical properties and forest floor nutrients under repeated prescribed-burning in eucalypt forests of south-east Queensland, Australia. New Zeal. J. For. Sci. 31, 170–187 (2001).

33. Glitzenstein, J. S., Streng, D. R. & Wade, D. D. Fire frequency effects on longleaf pine (Pinus palustris P. Miller) vegetation in South Carolina and northeast Florida, USA. Nat. Areas Journal. 23 22–37. 2003. (2003).

34. Le Quéré, C., Raupach, M. R., Canadell, J. G. & Marland, G. Trends in the sources and sinks of carbon dioxide. Nat. Geosci. 2, 831–836 (2009).

35. Bond, W. J., Woodward, F. I. & Midgley, G. F. The global distribution of ecosystems in a world without fire. New Phytol. 165, 525–538 (2005).

36. Jackson, R. B. et al. Trading water for carbon with biological carbon sequestration. Science (80-.). 310, 1944–1947 (2005).

37. Whitman, E., Parisien, M. A., Thompson, D. K. & Flannigan, M. D. Short-interval wildfire and drought overwhelm boreal forest resilience. Sci. Rep. 9, 1–12 (2019).

38. Hart, S. J. et al. Examining forest resilience to changing fire frequency in a fire-prone region of boreal forest. Glob. Chang. Biol. 25, 869–884 (2019).

39. Stephens, S. L. et al. Managing forests and fire in changing climates. Science vol. 342 41–42 (2013).

40. Steel, Z. L., Safford, H. D. & Viers, J. H. The fire frequency-severity relationship and the legacy of fire suppression in California forests. Ecosphere 6, 1–23 (2015).

41. Liu, Y. Y. et al. Recent reversal in loss of global terrestrial biomass. Nat. Clim. Chang. 5, 470–474 (2015).

42. Brandt, M. et al. Satellite passive microwaves reveal recent climate-induced carbon losses in African drylands. Nat. Ecol. Evol. 2, 827–835 (2018).

43. Pellegrini, A. F. A. et al. Fire frequency drives decadal changes in soil carbon and nitrogen and ecosystem productivity. Nature 553, 194–198 (2018).

44. Ma, Z. et al. Evolutionary history resolves global organization of root functional traits. Nature 555, 94–97 (2018).

45. Shah, F. et al. Ectomycorrhizal fungi decompose soil organic matter using oxidative mechanisms adapted from saprotrophic ancestors. New Phytol. 209, 1705–1719 (2016).

46. Averill, C., Bhatnagar, J. M., Dietze, M. C., Pearse, W. D. & Kivlin, S. N. Global imprint of mycorrhizal fungi on whole-plant nutrient economics. Proc. Natl. Acad. Sci. U. S. A. (2019) doi: 10.1073/pnas.1906655116.

47. Woinarski, J. C. Z., Risler, J. & Kean, L. Response of vegetation and vertebrate fauna to 23 years of fire exclusion in a tropical Eucalyptus open forest, Northern Territory, Australia. Austral Ecol. 29, 156–176 (2004).

48. Veenendaal, E. M. et al. On the relationship between fire regime and vegetation structure in the tropics. New Phytol. 218, 153–166 (2018).

49. Pellegrini, A. F. A. et al. Repeated fire shifts carbon and nitrogen cycling by changing plant inputs and soil decomposition across ecosystems. Ecol. Monogr. (2020) doi:10.1111/1365-2745.13351.

50. Johnson, D. W. & Curtis, P. S. Effects of forest management on soil C and N storage: meta analysis. For. Ecol. Manage. 140, 227–238 (2001).

51. Hijmans, R. J., Cameron, S. E., Parra, J. L., Jones, P. G. & Jarvis, A. Very high resolution interpolated climate surfaces for global land areas. Int. J. Climatol. 25, 1965–1978 (2005).

52. Harrison, X. A. et al. A brief introduction to mixed effects modelling and multi-model inference in ecology. PeerJ 2018, (2018).

53. Hedges, L. V., Gurevitch, J. & Curtis, P. S. The meta-analysis of response ratios in experimental ecology. Ecology 80, 1150–1156 (1999).

54. Gurevitch, J., Morrow, L. L., Wallace, A. & Walsh, J. S. A meta-analysis of competition in field experiments. Am. Nat. 140, 539–572 (1992).

55. Bates, D., Mächler, M., Bolker, B. & Walker, S. Fitting Linear Mixed-Effects Models using lme4. J. Stat. Softw. 67, 1–48 (2015).

56. Jackson, J. F., Adams, D. C. & Jackson, U. B. Allometry of constitutive defense: A model and a comparative test with tree bark and fire regime. Am. Nat. 153, 614–632 (1999).

57. Chave, J. et al. Towards a worldwide wood economics spectrum. Ecol Lett 12, 351–366 (2009).

58. Hoffmann, W. A., Marchin, R. M., Abit, P. & Lau, O. L. Hydraulic failure and tree dieback are associated with high wood density in a temperate forest under extreme drought. Glob. Chang. Biol. 17, 2731–2742 (2011).

59. Harmon, M. E. Decomposition of standing dead trees in the southern Appalachian Mountains. Oecologia 52, 214–215 (1982).

60. Zanne, A. E. et al. Three keys to the radiation of angiosperms into freezing environments. Nature 506, 89–92 (2014).

61. Pearse, W. D. et al. pez: phylogenetics for the environmental sciences. Bioinformatics 31, 2888–2890 (2015).

62. Kembel, S. W. et al. Picante: R tools for integrating phylogenies and ecology. Bioinformatics 26, 1463–1464 (2010).

63. Brockway, D. G. & Lewis, C. E. Long-term effects of dormant-season prescribed fire on plant community diversity, structure and productivity in a longleaf pine wiregrass ecosystem. For. Ecol. Manage. 96, 167–183 (1997).

64. Lewis, T. & Debuse, V. J. Resilience of a eucalypt forest woody understorey to long-term (34 - 55 years) repeated burning in subtropical Australia. Int. J. Wildl. Fire 21, 980–991 (2012).

65. Scudieri, C. A., Sieg, C. H., Haase, S. M., Thode, A. E. & Sackett, S. S. Understory vegetation response after 30 years of interval prescribed burning in two ponderosa pine sites in northern Arizona, USA. For. Ecol. Manage. 260, 2134–2142 (2010).

66. Lewis, T., Reif, M., Prendergast, E. & Tran, C. The effect of long-term repeated burning and fire exclusion on above- and below-ground Blackbutt (Eucalyptus pilularis) forest vegetation assemblages. Austral Ecol. 37, 767–778 (2012).

67. Stratton, R. Effects of long-term late winter prescribed fire on forest stand dynamics, small mammal populations, and habitat demographics in a Tennessee oak barrens. Masters Theses (2007).

68. Wade, D. D. Long-term site responses to season and interval of underburns on the Georgia Piedmont. For. Serv. Res. Data Arch. (2016) doi:10.2737/RDS-2016-0028.

69. Pellegrini, A. F. A., Hoffmann, W. A. & Franco, A. C. Carbon accumulation and nitrogen pool recovery during transitions from savanna to forest in central Brazil. Ecology 95, 342–352 (2014).

70. Nesmith, C. B., Caprio, A. C., Pfaff, A. H., McGinnis, T. W. & Keeley, J. E. A comparison of effects from prescribed fires and wildfires managed for resource objectives in Sequoia and Kings Canyon National Parks. For. Ecol. Manage. 261, 1275–1282 (2011).

71. Haywood, J. D., Harris, F. L., Grelen, H. E. & Pearson, H. A. Vegetative response to 37 years of seasonal burning on a Louisiana longleaf pine site. South. J. Appl. For. 25, 122–130 (2001).

72. Gignoux, J., Lahoreau, G., Julliard, R. & Barot, S. Establishment and early persistence of tree seedlings in an annually burned savanna. J. Ecol. 97, 484–495 (2009).

73. Tizon, F. R., Pelaez, D. V. & Elia, O. R. The influence of controlled fires on a plant community in the south of the Caldenal and its relationship with a regional state and transition model. Int. J. Exp. Bot. 79, 141–146 (2010).

74. Neill, C., Patterson, W. A. & Crary, D. W. Responses of soil carbon, nitrogen and cations to the frequency and seasonality of prescribed burning in a Cape Cod oak-pine forest. For. Ecol. Manage. 250, 234–243 (2007).

75. Ryan, C. M., Williams, M. & Grace, J. Above-and belowground carbon stocks in a miombo woodland landscape of Mozambique. Biotropica 43, 423–432 (2011).

76. Scharenbroch, B. C., Nix, B., Jacobs, K. A. & Bowles, M. L. Two decades of low-severity prescribed fire increases soil nutrient availability in a Midwestern, USA oak (Quercus) forest. Geoderma 183–184, 80–91 (2012).

77. Williams, R. J., Hallgren, S. W. & Wilson, G. W. T. Frequency of prescribed burning in an upland oak forest determines soil and litter properties and alters the soil microbial community. For. Ecol. Manage. 265, 241–247 (2012).

78. Stewart, J. F., Will, R. E., Robertson, K. M. & Nelson, C. D. Frequent fire protects shortleaf pine (Pinus echinata) from introgression by loblolly pine (P. taeda). Conserv. Genet. 16, 491–495 (2015).

79. Knapp, B. O., Stephan, K. & Hubbart, J. A. Structure and composition of an oak-hickory forest after over 60 years of repeated prescribed burning in Missouri, U.S.A. For. Ecol. Manage. 344, 95–109 (2015).

80. Olson, M. G. Tree regeneration in oak-pine stands with and without prescribed fire in the New Jersey Pine Barrens: management implications. North. J. Appl. For. 28, 47–49 (2011).

81. Whittaker, R. H. Gradient analysis of vegetation. Biol. Rev. Camb. Philos. Soc. 42, 207–264 (1967).

82. Bond, W. & Midgley, J. Ecology of sprouting in woody plants: the persistence niche. Trends Ecol. Evol. (Personal Ed. 16, 45–51 (2001).

